# EHRMENEIA: A Benchmark for Foundation Models on Longitudinal Electronic Health Records

**DOI:** 10.64898/2026.11.27.769542

**Authors:** Joana Gelabert-Xirinachs, Sergio Gosálvez, Blanca Alaejos-Pardo, Jordi Fonollosa, Alexandre Perera-Lluna

## Abstract

Electronic health record foundation models are evaluated using heterogeneous targets, vocabularies, and adaptation procedures, preventing systematic characterization across model families. We introduce EHRMENEIA (https://ehrmeneia.upc.edu/), a benchmark that provides a multidimensional evaluation framework for next-event prediction, multi-horizon event occurrence, trajectory fidelity and few-shot representation transfer. EHRMENEIA preserves each model’s native vocabulary and temporal representation while mapping clinical outputs to shared evaluation objects. Applied to two released clinical checkpoints and two synthetic demonstrations, the framework exposed substantial differences not only in performance but in the clinical questions that could be evaluated from each model–dataset interface. EHRMENEIA introduces a view of benchmarking in which the supported clinical evaluation domain is identified before performance within that domain is interpreted. The framework therefore shifts foundation-model benchmarking from scalar ranking toward characterization of what can, and cannot, be inferred from model outputs.

---

Foundation models (FMs) are deep learning models pretrained on broad datasets, typically through self-supervised learning, and designed to support adaptation across multiple downstream tasks [1]. In medicine, FMs are increasingly used to learn from large-scale biomedical and clinical data, with applications ranging from clinical-event prediction and medication recommendation to the study of disease onset, comorbidity, progression and healthcare resource management [2, 3]. Their evaluation, however, depends fundamentally on the type of information from which they learn.

Medical FMs can be broadly distinguished by whether pretraining is driven primarily by secondary biomedical corpora or by primary data generated during clinical care. Models trained on scientific literature, guidelines, textbooks and related textual sources learn representations of biomedical knowledge expressed through language [4, 5, 6, 7, 8, 9]. Their evaluation accordingly focuses on language-mediated capabilities such as examination-style question answering [10, 11], medical dialogue [12], instruction alignment [13], clinical documentation [14], healthcare agents [15] and reasoning over EHR-derived context [16]. These benchmarks address distinct and non-trivial evaluation problems, but their object of evaluation is primarily the model’s behavior when biomedical or clinical information is expressed through language.

Longitudinal EHR foundation models learn from a different observational object. Diagnoses, medications, laboratory measurements, procedures, and other records are collected in clinical care settings and form temporally ordered patient trajectories. EHR foundation models have evolved from BERT-style masked-event representation learning toward autoregressive sequence modeling [17, 18, 19, 20], enabling them not only to encode patient histories for downstream prediction but also to forecast subsequent clinical events and generate future trajectories. Their evaluation must therefore interrogate properties of a longitudinal clinical process: which events are predicted, when they occur, how generated trajectories evolve over time and what information remains accessible in the learned representations.

This distinction changes the evaluation problem. For longitudinal foundation models, the central question is not simply whether a model performs well on a predefined task, but what constitutes evidence that it has learned clinically meaningful temporal structure. This question has become increasingly important as foundation-model benchmarking moves beyond estimating performance on isolated predictive tasks toward determining which properties of broadly pretrained models can actually be tested. Recent work has argued that benchmarking biomedical foundation models requires the community to define not only metrics but also the questions, limitations and forms of evidence that an evaluation is intended to probe [21]. Existing longitudinal EHR evaluations span next-event forecasting, downstream prediction, representation transfer and synthetic-trajectory generation, but these are typically assessed separately and under different targets, horizons, coding systems and adaptation procedures (see Table 1).

**Table 1:**
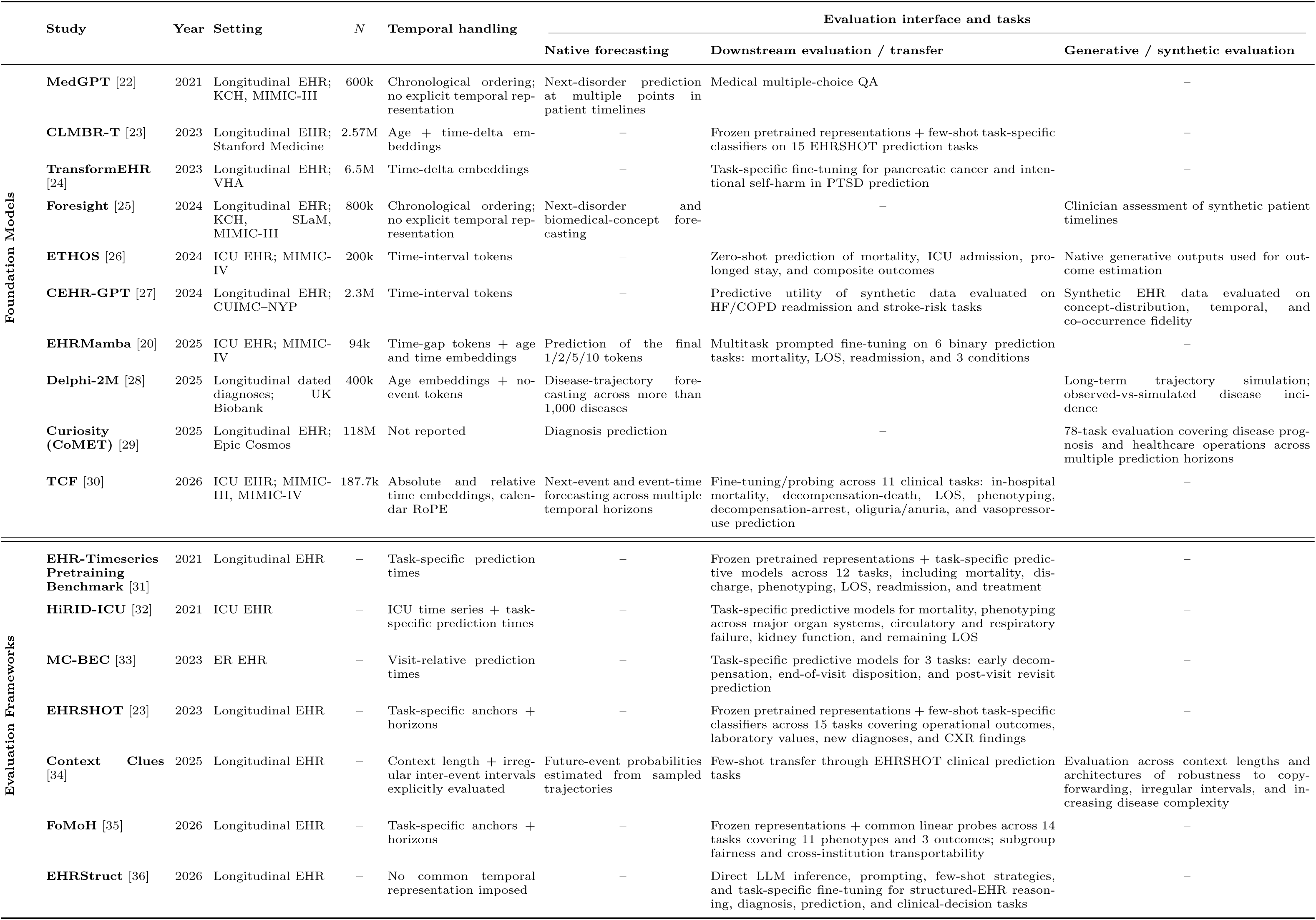
Evaluation landscape for longitudinal EHR foundation models and evaluation frameworks. Foundation-model studies and evaluation frameworks are compared according to data setting, reported pretraining population size (*N* ), temporal handling, and evaluation interface. For foundation models, temporal handling refers to the model-native representation of time; for evaluation frameworks, it refers to temporal information required or explicitly evaluated by the protocol.

|  | Study | Year | Setting | *N* | Temporal handling | Evaluation interface and tasks |  |  |
|---|---|---|---|---|---|---|---|---|
|  |  |  |  |  |  | Native forecasting | Downstream evaluation / transfer | Generative / synthetic evaluation |
| Foundation Models | **MedGPT** [22] | 2021 | Longitudinal EHR; KCH, MIMIC-III | 600k | Chronological ordering; no explicit temporal rep-resentation | Next-disorder prediction at multiple points in patient timelines | Medical multiple-choice QA | – |
|  | **CLMBR-T** [23] | 2023 | Longitudinal EHR; Stanford Medicine | 2.57M | Age + time-delta em-beddings | – | Frozen pretrained representations + few-shot task-specific classifiers on 15 EHRSHOT prediction tasks | – |
|  | **TransformEHR** [24] | 2023 | Longitudinal EHR; VHA | 6.5M | Time-delta embeddings | – | Task-specific fine-tuning for pancreatic cancer and inten-tional self-harm in PTSD prediction | – |
|  | **Foresight** [25] | 2024 | Longitudinal EHR; KCH, SLaM, MIMIC-III | 800k | Chronological ordering; no explicit temporal rep-resentation | Next-disorder and biomedical-concept fore-casting | – | Clinician assessment of synthetic patient timelines |
|  | **ETHOS** [26] | 2024 | ICU EHR; MIMIC-IV | 200k | Time-interval tokens | – | Zero-shot prediction of mortality, ICU admission, pro-longed stay, and composite outcomes | Native generative outputs used for out-come estimation |
|  | **CEHR-GPT** [27] | 2024 | Longitudinal EHR; CUIMC–NYP | 2.3M | Time-interval tokens | – | Predictive utility of synthetic data evaluated on HF/COPD readmission and stroke-risk tasks | Synthetic EHR data evaluated on concept-distribution, temporal, and co-occurrence fidelity |
|  | **EHRMamba** [20] | 2025 | ICU EHR; MIMIC-IV | 94k | Time-gap tokens + age and time embeddings | Prediction of the final 1/2/5/10 tokens | Multitask prompted fine-tuning on 6 binary prediction tasks: mortality, LOS, readmission, and 3 conditions | – |
|  | **Delphi-2M** [28] | 2025 | Longitudinal dated diagnoses; UK Biobank | 400k | Age embeddings + no-event tokens | Disease-trajectory fore-casting across more than 1,000 diseases | – | Long-term trajectory simulation; observed-vs-simulated disease inci-dence |
|  | **Curiosity (CoMET)** [29] | 2025 | Longitudinal EHR; Epic Cosmos | 118M | Not reported | Diagnosis prediction | – | 78-task evaluation covering disease prog-nosis and healthcare operations across multiple prediction horizons |
|  | **TCF** [30] | 2026 | ICU EHR; MIMIC-III, MIMIC-IV | 187.7k | Absolute and relative time embeddings, calen-dar RoPE | Next-event and event-time forecasting across multiple temporal horizons | Fine-tuning/probing across 11 clinical tasks: in-hospital mortality, decompensation-death, LOS, phenotyping, decompensation-arrest, oliguria/anuria, and vasopressor-use prediction | – |
| Evaluation Frameworks | **EHR-Timeseries Pretraining Benchmark** [31] | 2021 | Longitudinal EHR | – | Task-specific prediction times | – | Frozen pretrained representations + task-specific predic-tive models across 12 tasks, including mortality, dis-charge, phenotyping, LOS, readmission, and treatment | – |
|  | **HiRID-ICU** [32] | 2021 | ICU EHR | – | ICU time series + task-specific prediction times | – | Task-specific predictive models for mortality, phenotyping across major organ systems, circulatory and respiratory failure, kidney function, and remaining LOS | – |
|  | **MC-BEC** [33] | 2023 | ER EHR | – | Visit-relative prediction times | – | Task-specific predictive models for 3 tasks: early decom-pensation, end-of-visit disposition, and post-visit revisit prediction | – |
|  | **EHRSHOT** [23] | 2023 | Longitudinal EHR | – | Task-specific anchors + horizons | – | Frozen pretrained representations + few-shot task-specific classifiers across 15 tasks covering operational outcomes, laboratory values, new diagnoses, and CXR findings | – |
|  | **Context Clues** [34] | 2025 | Longitudinal EHR | – | Context length + irreg-ular inter-event intervals explicitly evaluated | Future-event probabilities estimated from sampled trajectories | Few-shot transfer through EHRSHOT clinical prediction tasks | Evaluation across context lengths and architectures of robustness to copy-forwarding, irregular intervals, and in-creasing disease complexity |
|  | **FoMoH** [35] | 2026 | Longitudinal EHR | – | Task-specific anchors + horizons | – | Frozen representations + common linear probes across 14 tasks covering 11 phenotypes and 3 outcomes; subgroup fairness and cross-institution transportability | – |
|  | **EHRStruct** [36] | 2026 | Longitudinal EHR | – | No common temporal representation imposed | – | Direct LLM inference, prompting, few-shot strategies, and task-specific fine-tuning for structured-EHR reason-ing, diagnosis, prediction, and clinical-decision tasks | – |

The challenge is therefore not only to standardize metrics, but to define the object to which those metrics refer. A patient’s underlying clinical state is not observed directly in an EHR. It becomes partially observable through interactions with the healthcare system and is subsequently documented, measured and coded before reaching a model. The same clinical phenomenon may therefore be represented through different coding systems, levels of granularity, temporal conventions and model-specific tokenization schemes. Comparing model-native outputs directly can consequently confound differences in learned distributions with differences in how clinical observations are represented. Evaluation must instead be defined at a level where clinically interpretable questions can be related consistently to observable model outputs.

We developed EHRMENEIA to define this evaluation locus. The benchmark distinguishes the model-native representation space from a shared clinical evaluation space, allowing models to retain their original vocabularies, tokenization strategies and representations of time while exposing clinically interpretable outputs to common evaluation questions. EHRMENEIA therefore standardizes neither the model nor its internal representation; it defines the clinical questions against which observable model behavior can be interrogated.

Importantly, standardization does not imply universal comparability. Whether a particular quality can be compared still depends on whether the evaluated model and dataset support the same outcome definitions and sufficient statistical support. EHRMENEIA therefore treats coverage, support and non-estimability as part of the benchmark result, allowing it to distinguish comparisons that are well defined from those that are only partial or unsupported.

Its name derives from the Greek *hermēneia*, meaning interpretation or explanation and associated with Aristotle’s *Peri Hermēneias*. It evokes the relation between things and the signs through which they become intelligible, reflecting the central problem addressed by the benchmark: determining which claims about a model remain interpretable across successive layers of observation, documentation, coding and computational representation.

EHRMENEIA evaluates complementary properties of longitudinal EHR foundation models across three dimensions and four evaluation levels (Fig. 1). Predictive Competence measures native next-event forecasting. Clinical and Population Fidelity assess whether the autoregressive distribution remains faithful when propagated across future time windows and complete generated trajectories. Representation Transferability evaluates whether information acquired during pretraining can be reused for previously unseen prediction tasks under controlled few-shot adaptation with a frozen backbone. To our knowledge, no existing benchmark jointly evaluates these dimensions within a common clinical evaluation space. Together, these levels define a prespecified space of complementary evaluation questions rather than a hierarchy culminating in a single score. Level 4 is deliberately extensible: its released task catalogue provides a reference set of transfer questions, while additional clinically defined anchor–target–horizon tasks can extend this region without altering the underlying evaluation protocol.

**Figure 1:**
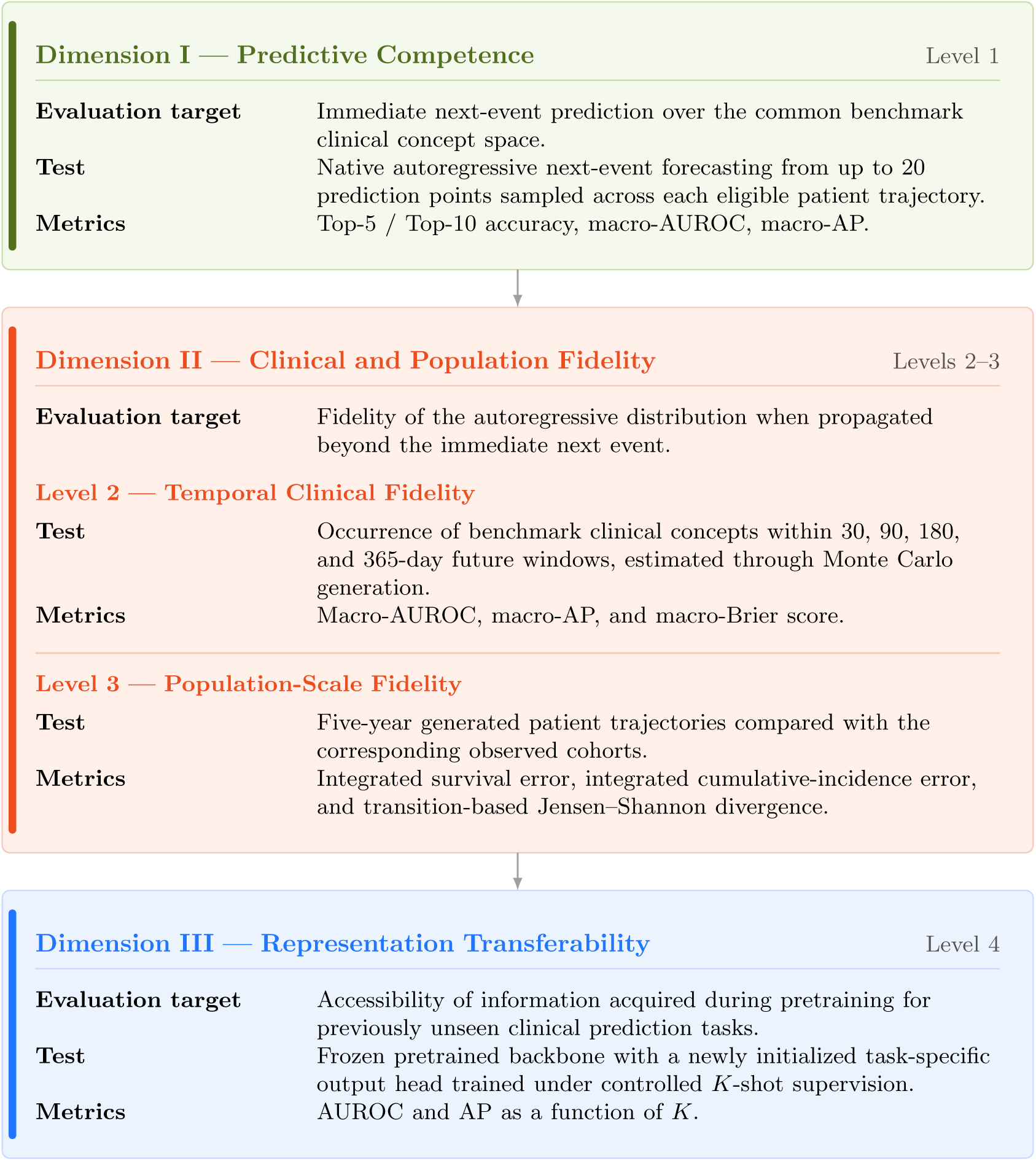
Multidimensional evaluation framework for EHR foundation models. Within each level, the figure summarizes the evaluation target, benchmark test, and primary reported metrics. Together, the levels define complementary regions of a common clinical evaluation space rather than components of a composite score.

We applied EHRMENEIA to four heterogeneous foundation-model implementations: ETHOS, CLMBR-T, Delphi, and CEHR-GPT (https://ehrmeneia.upc.edu/). These applications test whether a common evaluation space can be instantiated across substantially different vocabularies, temporal representations and model interfaces, and whether each implementation can be characterized by both the region of that space it supports and its measured behavior within that region.

## Results

We applied EHRMENEIA to four heterogeneous implementations, comprising two released clinical checkpoints (CLMBR-T on EHRSHOT and ETHOS on MIMIC-IV 2.2 [37]) and two locally trained synthetic demonstrations (Delphi and CEHR-GPT). Among the models summarized in Table 1, we selected implementations for which the combination of released model interfaces, executable code and accessible evaluation data allowed the EHRMENEIA workflow to be instantiated. Implementation details can be found both in Section 1 of the Supplementary Materials and the shared repository.

Across implementation-specific evaluation cohorts, the benchmark processed 359,037 next-event prediction points from 33,098 patients, 2,151,600 generated futures and 2,320 frozen-backbone adaptation heads. These applications were selected to examine whether a common evaluation protocol could accommodate different model interfaces, vocabularies and temporal representations, rather than to establish a ranking of model families. Figure 2 summarizes their evaluable domains and performance profiles, while Figure 3 presents the underlying level-specific results. Complete numerical results are reported in the Supplementary Tables and on the benchmark website.

**Figure 2:**
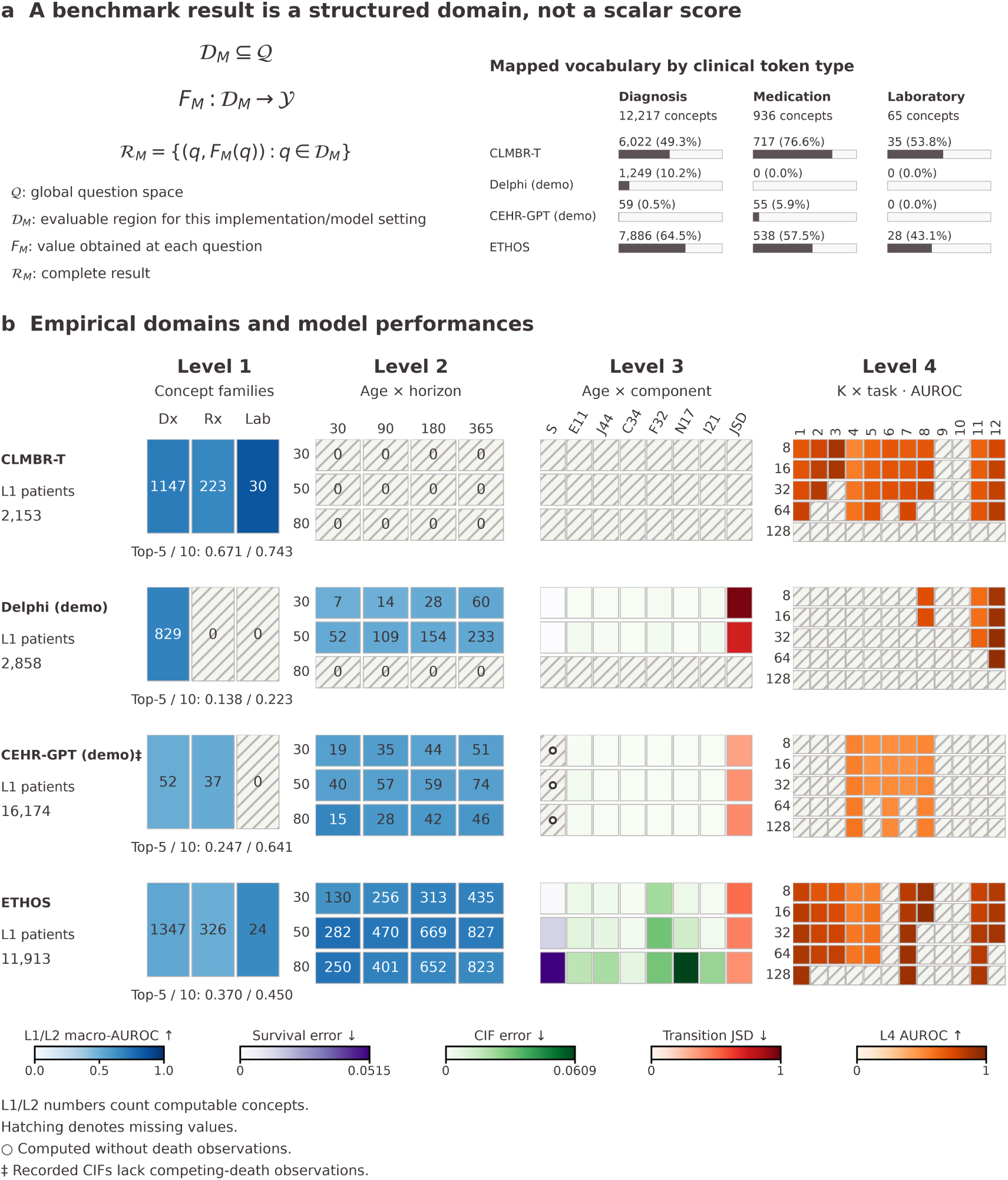
EHRMENEIA characterizes model performance over a structured domain of evaluable questions. **a,** The benchmark question space *Q* contains an evaluable subset *D_M_* for each model implementation and evaluation setting. The mapping *F_M_* assigns a metric value to each evaluable question *q*; *R_M_* collects the resulting question–value pairs. Vocabulary coverage is shown separately for diagnoses, medications and laboratory concepts. **b,** Evaluation profiles for CLMBR-T, Delphi, CEHR-GPT and ETHOS. Delphi and CEHR-GPT are locally trained synthetic demonstrations. Level 1 cells represent cl_7_inical domains; Level 2 cells represent index-age and prediction-horizon combinations. In both levels, colour encodes macro-AUROC and numbers indicate computable concepts. Level 1 also reports Top-5 and Top-10 accuracy. Level 3 shows integrated absolute survival error, integrated absolute cumulative-incidence error for six ICD-10 diagnosis families, and transition Jensen–Shannon divergence (JSD). Level 4 shows mean AUROC across 20 support draws for each task and *K* setting.

**Figure 3:**
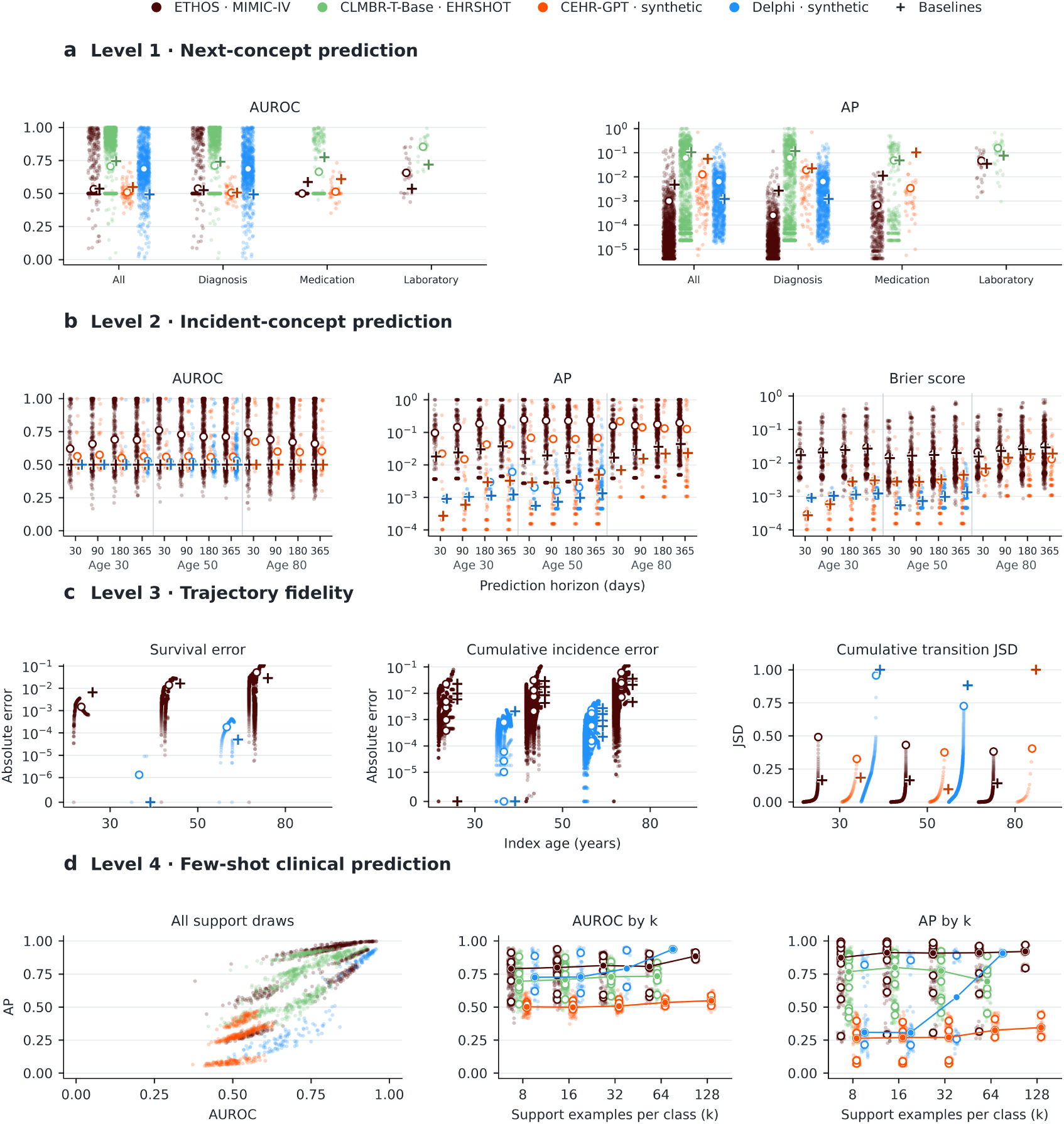
EHRMENEIA benchmark results across four evaluation levels. **a,** AUROC and average precision (AP) for next-concept prediction by concept domain and overall. Crosses mark the personal history-frequency baseline, which divides each mapped concept count by the total number of clinical events retained in the model input. **b,** AUROC, AP and Brier score for incident-concept prediction by index age and prediction horizon. Crosses mark the marginal-prevalence baseline estimated in a patient-disjoint adaptation cohort. Small points in **a** and **b** represent individual concepts, and open circles indicate macro-averaged scores. **c,** Survival and cumulative incidence errors by index age. Small points show absolute differences between observed and generated curves at published follow-up times, and open circles show integrated errors. For transition JSD, points show the running sum of pairwise contributions _8_in ascending order, with the open circle marking the total. Crosses mark the median observed–observed discrepancy across valid patient-disjoint split-half comparisons. CEHR-GPT is omitted from the survival and cumulative incidence panels because its evaluation data lack death observations. **d,** AUROC and AP for each estimable support draw, shown jointly and by the number of support examples per class (k). Open circles mark task means, and lines connect the median of the estimable task means at each *k* for each model. Evaluation cohorts and estimable tasks vary across models.

### Models occupied different evaluation domains

EHRMENEIA projected eligible model outputs onto a shared space of 13,218 clinical concepts, comprising 12,217 diagnoses, 936 medication classes and 65 laboratory concepts. The mapping preserved clinically defined concept identities without requiring models to share a native vocabulary, tokenization scheme or temporal representation. Ambiguous mappings were excluded, and broader native codes were not treated as evidence for narrower benchmark concepts. The released concept space and model-specific terminology mappings are available in the benchmark repository and website.

Despite this common semantic reference, the supported evaluation domains differed substantially. CLMBR-T exposed next-event predictions and pretrained representations but lacked a validated timed-trajectory generation interface; Delphi’s demonstration supported diagnosis-only outputs; and CEHR-GPT lacked laboratory and mortality observations. Consequently, not every implementation could be evaluated at every level or for every clinical outcome. These differences were not treated as missing values around a common score: they defined the set of clinical claims that could be supported by each model–dataset pair.

### Common metrics did not imply comparable clinical questions

Level 1 illustrated how a shared metric can summarize different empirical evaluation sets. Next-event forecasting yielded 1,400, 1,697, 829 and 89 computable concepts for CLMBR-T, ETHOS, Delphi and CEHR-GPT, respectively. Thus, macro-AUROC and macro-AP were calculated over markedly different clinical outcome panels, limiting the interpretation of direct numerical comparisons between implementations.

Metric choice introduced a further distinction. The relative ordering of implementations changed between macro-AUROC, Top-5 and Top-10 accuracy, reflecting differences between concept-level discrimination and within-query ranking. The patient-history frequency reference also showed that performance relative to a simple predictor depended on the metric and clinical domain (Supplementary Table **??**). Together, these findings demonstrate that a common scoring procedure does not, by itself, establish a common clinical estimand or a universal ordering of model quality.

### Complementary fidelity measures revealed distinct model properties

Levels 2 and 3 evaluated properties of longitudinal behavior that cannot be inferred from next-event performance alone. Level 2 assessed event occurrence within predefined future windows, while Level 3 compared generated and observed trajectories through population-level survival, cumulative incidence and local event transitions.

In Level 2, ETHOS achieved higher macro-AUROC and macro-AP than the marginal-prevalence reference across all 12 evaluable age–horizon settings, whereas the reference consistently yielded lower macro-Brier scores (Supplementary Table **??**). Discrimination and probability prediction error therefore conveyed different information about the same generated-future probabilities.

Level 3 revealed a related distinction between marginal and sequential fidelity. For ETHOS, relatively small survival and cumulative-incidence discrepancies coexisted with transition Jensen– Shannon divergence (JSD) values of 0.38–0.49. Observed–observed split-half comparisons helped contextualize these distances: at age 50, transition JSD was 0.431 for ETHOS and 0.727 for the Delphi demonstration, whereas the corresponding observed–observed medians were 0.164 and 0.882 (Supplementary Table **??**). These contrasts depend on the empirical support of the observed transition distributions and do not establish a universal fidelity threshold. Moreover, CEHR-GPT’s missing mortality observations prevented clinically valid evaluation of survival and competing-risk incidence, regardless of whether numerical summaries could be computed. Together, these results show why marginal agreement, sequential fidelity and the conditions supporting their estimation must be assessed separately.

### Representation transfer depended on task and empirical support

Level 4 extended the evaluation beyond native prediction and generation to the information accessible in frozen pretrained representations. Few-shot adaptation curves differed across implementations and clinical tasks. ETHOS showed gains with increasing supervision on several outcomes, including HbA1c-to-diabetes, kidney disease and cardiovascular prediction, whereas CLMBR-T exhibited task-dependent improvements and several synthetic-demonstration settings remained close to chance or were not estimable.

Importantly, non-estimability also arose from insufficient positive or negative outcome support, even when a model exposed the required representation interface. Thus, the extent of the evaluable transfer domain depended jointly on model capabilities, available observations and the support required by the adaptation protocol.

### Benchmark output is a domain, not a score

Taken together, these results indicate that benchmark output cannot be represented adequately by a single scalar. Let *Q* denote the prespecified space of clinical evaluation questions. For a given model–dataset implementation (M), the evaluable domain is

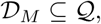

and performance defines a mapping

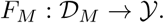

The benchmark result is therefore characterized jointly by the domain of supported questions and the metric field defined over that domain. In this sense, a benchmark result is a region, not a point.

### EHRMENEIA supported decentralized evaluation

All four applications were executed locally using their native model implementations and evaluation data. EHRMENEIA then reduced the resulting outputs to standardized aggregate metrics, support information and metadata, automatically assembling portable benchmark packages for visualization and submission. This separation enabled a common evaluation workflow without requiring patient-level records, predictions, learned representations or generated trajectories to be transferred from the local evaluation environment.

## Discussion

To our knowledge, EHRMENEIA is the first framework to evaluate longitudinal EHR foundation models across native forecasting, temporal occurrence, population-scale trajectory fidelity and representation transfer within a common clinical evaluation space. Rather than treating benchmarking as the assignment of scores to models, EHRMENEIA defines a set of clinically interpretable evaluation questions and determines which of them can be interrogated through a given model–dataset interface. The four empirical applications show that this abstraction can operate across heterogeneous implementations while preserving their native architectures, vocabularies and temporal representations.

A second contribution is to make comparability an explicit property of the evaluation rather than an implicit assumption. Terminology coverage, available outcomes, temporal generation capabilities and statistical support determine which quantities are estimable for a given implementation. For each model *M* , EHRMENEIA first identifies the supported subset *D_M_* of a prespecified evaluation space and only then characterizes behavior within that subset. In this sense, the benchmark output is defined by its evaluable domain: non-estimability is informative because it identifies questions that the available interface, semantic mapping or observations do not justify answering.

This formulation also explains why longitudinal foundation-model evaluation should not be collapsed into a single scalar. The empirical applications showed that the evaluation dimensions interrogate distinct properties of model behavior. In particular, agreement in marginal survival or recorded disease-incidence curves did not imply agreement in local sequential event structure, and increasing supervision did not uniformly recover transferable signal across tasks and implementations. The observed–observed references further illustrated that the interpretation of trajectory-level discrepancies depends on the empirical variability and support of the evaluated cohort. Longitudinal model quality therefore cannot be inferred from any single projection of model behavior, nor can a metric value be interpreted independently of the conditions under which it was estimated. A further design principle is that EHRMENEIA standardizes the evidence exposed for evaluation rather than the model itself. Model execution, tokenization, temporal representation and generation remain native to each implementation; only clinically interpretable outputs are mapped to common evaluation objects. This separation also enables decentralized evaluation. Models and data can remain within their local environment, while the evaluation pipeline reduces the resulting outputs to standardized aggregate metrics, support information and metadata. In the released implementation, these outputs are automatically assembled into a portable benchmark package for visualization and submission, without requiring patient-level records, predictions, representations or generated trajectories to leave the local environment. This provides a practical route for applying a common benchmark across institutions where direct pooling of clinical data or model outputs may be infeasible.

However, the present validation has several limitations. Two applications used released clinical checkpoints, whereas the Delphi and CEHR-GPT experiments were synthetic implementation demonstrations of the original models. Broader evaluation across additional checkpoints, institutions and datasets will therefore be necessary to determine how the resulting evaluation domains generalize. The released EHRMENEIA space is also not exhaustive: its four levels and current task catalogue provide an initial operationalization of clinically meaningful model properties that does not aim to be understood as a definitive ontology of longitudinal foundation-model capabilities. Finally, all results must be interpreted in relation to the observational process that generates EHR data. EHRMENEIA evaluates fidelity to documented healthcare trajectories, which are shaped by clinical practice, coding, measurement frequency, access to care and institutional workflows as well as by underlying patient health.

EHRMENEIA should therefore be viewed both as a benchmark and as an initial map of the evaluation space for longitudinal EHR foundation models. As clinical foundation models become increasingly general, the methodological question shifts from how these models should be scored to what can be concluded from their observable predictions, representations and generated trajectories. EHRMENEIA provides one operational answer by defining a domain of supported clinical questions together with the corresponding metric field. Extending this map will require community agreement on which additional regions of model behavior constitute meaningful evidence. The aim is not a universal leaderboard, but a shared map and compass for navigating what longitudinal foundation models can validly be said to have learned.

## Online Methods

### Benchmark representation and terminology

EHRMENEIA represents each eligible clinical event as

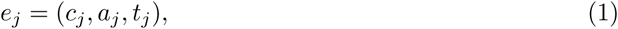

where *c_j_* is the benchmark clinical concept, *a_j_* contains optional event attributes, and *t_j_* is event time. The common concept space *V_B_* contains diagnoses represented at up to four-character ICD-10 [38] level, medications at ATC [39] level 4 and laboratory-test identities. Numerical laboratory values are retained as optional attributes rather than represented as distinct benchmark concepts.

Models retain their native vocabularies, tokenization schemes and representations of time. Cross-model evaluation is performed only after eligible native outputs have been projected onto *V_B_*. When multiple native outputs map unambiguously to the same benchmark concept, their predictive mass is combined. Outputs without an unambiguous mapping are not assigned to narrower benchmark concepts. Mapping coverage is retained as part of the benchmark output.

For generative evaluation, future trajectories are produced using each model’s native generation procedure and subsequently converted to benchmark events with an interpretable elapsed time from the prediction index. EHRMENEIA therefore standardizes the clinical evaluation object rather than the internal representation or generation mechanism of the model.

Contextual information is an important component of longitudinal EHR modeling because the interpretation and predictive relevance of a clinical event may depend on demographic, temporal, administrative, or other aspects of the observation context. We denote by **x***_i_*(*t*) the contextual information available for patient *i* up to time *t*. Such information may be used by the model together with the observed clinical-event history when estimating future events. However, EHRMENEIA does not evaluate contextual variables as prediction targets themselves. Instead, their potential value is reflected indirectly through the predictive and generative behavior of models that condition on them.

### Common evaluation setting

Developers must declare and ensure patient-level disjointness between the data used to pretrain the submitted model and all data used for EHRMENEIA evaluation, including adaptation, support and test cohorts.

Indexing is defined separately according to the evaluation target. Level 1 does not use fixed age anchors. Levels 2 and 3 retain standardized reference ages of 30, 50, and 80 years. At these reference ages, only events recorded before the index time are available to the model. The reference ages were chosen to sample clinically distinct stages of the adult life course, spanning early adulthood, midlife and older age, where disease burden, multimorbidity and the density of recorded healthcare observations differ substantially.

For Levels 2 and 3, patients contribute only at reference ages for which the required history and follow-up can be defined. The same patient may therefore contribute at multiple reference ages, with each eligible patient–index-age pair treated as a separate longitudinal evaluation case. Cohort sizes and support are reported separately for every evaluation condition. The reported summaries do not assume statistical independence among cases contributed by the same individual, and no hypothesis testing based on case-level independence is performed.

### Dimension I: Predictive Competence

#### Level 1: Native next-event prediction

In Level 1, each candidate prediction point must have a preceding recorded history and a subsequent eligible benchmark concept available. Then, the interval between the earliest and latest eligible candidate points is divided into up to 20 equal-duration intervals. One candidate is sampled from each occupied interval; when fewer than 20 intervals contain candidates, the remaining prediction points are sampled without replacement from the unused eligible candidates until either 20 points have been selected or no candidates remain. A target record can contribute to at most one selected prediction point.

Top-5 and Top-10 accuracy evaluate whether the target is among the five or ten highest-ranked eligible benchmark concepts. To account for differences in the number of prediction points contributed by each patient, each selected point is assigned a weight of 1*/T_i_*, where *T_i_* denotes the number of selected prediction points for patient *i* within the evaluation stratum. One-versus-rest AUROC and average precision (AP) are calculated for each benchmark concept with both positive and negative observations and macro-averaged across the computable-concept panel. Concept-level positive and negative patient counts and weighted outcome prevalence are retained to characterize the empirical support of each estimate. Because outcomes are defined at individual prediction points, the same patient may contribute both positive and negative observations for a given concept.

As a reference, we used the relative frequency of each benchmark concept in the patient’s history preceding each prediction point.

### Dimension II: Clinical and Population Fidelity

#### Level 2: Temporal Clinical Fidelity

For each index time *t_i_*, evaluation is performed over a set of prediction horizons Δ*_r_ ∈ D*, where *D* denotes the set of evaluation horizons.

In the reference EHRMENEIA configuration, *D* = *{*30, 90, 180, 365*}* days. Alternative horizon sets may be defined when adapting the benchmark to clinical settings characterized by different temporal scales.

For patient *i*, benchmark concept *v*, and horizon Δ, the observed outcome is

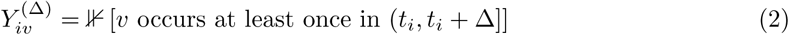

For an autoregressive model, the corresponding occurrence probability generally requires marginalization over all possible future continuations and is therefore not directly available. The benchmark approximates it through Monte Carlo generation. For each observed history, *M* independent future trajectories are generated. Let *G_im_* denote the *m*-th generated future trajectory for patient *i*, after generated events have been mapped to benchmark concepts and represented by their elapsed time from the index.

The event-occurrence probability is estimated as

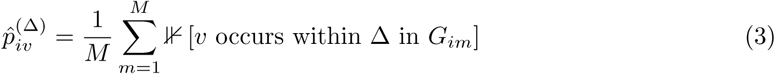

Thus, 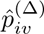 is the proportion of sampled futures in which concept *v* occurs within the specified horizon. This definition is independent of the model’s internal representation of time, provided that generated outputs can be transformed into benchmark concepts with interpretable elapsed times from the prediction index.

The Monte Carlo estimate is used as a patient-level event-occurrence probability. Within each index-age and prediction-horizon setting, discrimination is evaluated using AUROC and AP, calculated for each concept with both observed outcome classes and macro-averaged across the computable-concept panel. Probability prediction quality is additionally assessed using the Brier score, defined for each concept as the mean squared difference between predicted occurrence probabilities and observed binary outcomes. Concept-level Brier scores are macro-averaged over the same computable-concept panel as AUROC and AP. Unlike discrimination metrics, the Brier score also reflects the numerical accuracy of the predicted probabilities, although it does not isolate calibration from other components of predictive performance. All metrics are evaluated on observations with determinate outcome status according to the follow-up eligibility criteria described below.

Because occurrence probabilities are estimated from a finite number of generated trajectories, their Monte Carlo standard error is approximated by

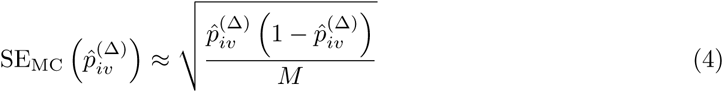

The reference configuration uses *M* = 100, which bounds the patient-level Monte Carlo standard error of the estimated occurrence probability by 0.05.

##### Observed follow-up and censoring

A positive outcome can be established whenever the target concept is observed within the prediction window. In contrast, assigning a negative outcome requires sufficient follow-up to establish that the event did not occur throughout the complete interval. Accordingly, a patient contributes to the negative class at (*t_i_,* Δ) only when the record remains observable through *t_i_*+ Δ.

Patients whose records end before completion of the window without experiencing the target event are treated as having indeterminate outcome status and are excluded from that concept– horizon evaluation. The number of eligible positive and negative patients is reported for each index-age, concept, and prediction-horizon combination. This rule avoids interpreting incomplete observation as evidence of event non-occurrence while making follow-up-based selection explicit.

We used marginal event prevalence as a reference, estimated separately for each concept, index age and horizon in a patient-disjoint adaptation cohort. The estimate was the proportion of positive outcomes among cases with determinate follow-up; settings without eligible adaptation outcomes received a probability of 0.5. All test patients in a given setting received the same estimated probability. Reference AUROC, AP and Brier scores were evaluated using the same test cases and computable-concept panels as the model.

#### Level 3: Population-Scale Fidelity

For each eligible patient *i*, the observed history *H_i_*(*t_i_*) is provided to the model and *M_G_* independent future trajectories are generated autoregressively up to a common evaluation horizon *τ* = 5 years. Generated events are projected onto the benchmark clinical concept space and represented relative to the index time. The same evaluation horizon [0*, τ* ] is applied to observed and generated trajectories.

Population-scale fidelity is evaluated along three complementary axes: survival, disease incidence, and sequential clinical-event structure.

##### Survival fidelity

Let 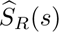 denote the survival function estimated from the observed cohort and 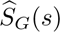 the corresponding function estimated from generated trajectories, where *s* denotes elapsed time from the index. Observed survival is estimated in the presence of right censoring, while generated trajectories that do not contain a death event before *τ* are administratively censored at the end of the generation horizon.

Native trajectory termination without an explicit death event is treated as the end of recorded follow-up and is not interpreted as death. Mortality-fidelity evaluation is restricted to settings in which death is represented in the observed data and can be identified in generated trajectories.

Agreement between observed and generated survival is quantified using the integrated survival error,

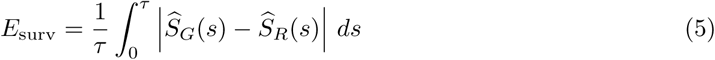

A value of zero indicates identical estimated survival functions over the evaluation interval, while increasing values indicate greater disagreement. Survival curves and numbers at risk are reported alongside the integrated error so that the temporal origin of discrepancies remains visible rather than being represented only by a scalar summary.

##### Disease incidence fidelity

Disease incidence fidelity is evaluated for a predefined set of clinical conditions selected to represent heterogeneous temporal and clinical dynamics: Type 2 Diabetes (`ICD-10: E11`), Chronic Obstructive Pulmonary Disease (`ICD-10: J44`), Lung Cancer (`ICD-10: C34`), Depressive Episodes (`ICD-10: F32`), Acute Kidney Injury (`ICD-10: N17`), and Acute Myocardial Infarction (`ICD-10: I21`). These targets span chronic, oncological, psychiatric, and acute conditions, providing complementary tests of whether generated trajectories reproduce disease incidence patterns across clinically distinct temporal profiles. For each condition, evaluation is restricted to patients without a recorded diagnosis belonging to the corresponding ICD-10 prefix family before the index time. The outcome is defined as the first recorded post-index occurrence of an eligible diagnosis in that family.

For condition *k*, let 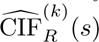 and 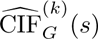 denote the observed and generated cumulative incidence functions for first occurrence of the condition after the index.

Death before onset of the target condition is treated as a competing event rather than as non-informative censoring. Cumulative incidence is therefore estimated using a competing-risk formulation, with Aalen–Johansen estimates used for both observed and generated cohorts.

Agreement is summarized by the integrated cumulative-incidence error,

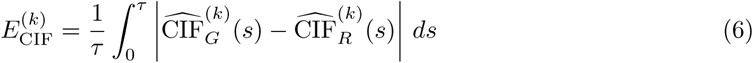

Lower values indicate closer agreement between observed and generated disease incidence trajectories. As for survival, cumulative-incidence curves are reported together with the integrated summary metric.

##### Sequential transition fidelity

Survival and disease incidence characterize clinically important temporal outcomes but do not determine whether generated trajectories reproduce clinically relevant sequential structure. Level 3 therefore additionally compares transitions between temporally consecutive benchmark events. Sequential transition fidelity is evaluated using a set of predefined diagnosis groups corresponding to the same conditions used for disease-incidence analysis.

For each occurrence of these chosen diagnoses, the benchmark records the concept or concepts (if there are co-occurring events) occurring at the next distinct event time. Thus, diagnoses define the source states, whereas destination states may correspond to any eligible benchmark concept, including diagnoses, medications, and laboratory events. Events recorded at the same timestamp are treated as unordered: no transitions are defined among them, while all eligible concepts occurring at the next distinct event time are considered outgoing transitions from the preceding diagnosis.

To characterize the empirical support of each transition distribution, the benchmark additionally reports the total number of observed and generated transitions and the number of distinct transition pairs in each population.

Transition distributions are defined over the union of diagnosis-to-next-event pairs observed in either the real or generated population. Let *q_R_* and *q_G_* denote the normalized distributions of these transitions in the observed and generated populations, respectively. Sequential fidelity is quantified using Jensen–Shannon divergence,

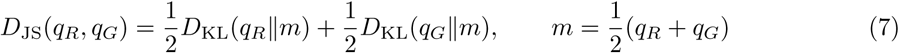

Using base-2 logarithms, *D*_JS_ ranges from 0 to 1, where lower values indicate closer agreement between observed and generated local event structure following the predefined conditions.

##### Sampling uncertainty and observed–observed references

For Levels 1–3, sampling uncertainty was characterized using a patient-level Bayesian bootstrap with 1,000 replicates. In each replicate, independent exponential weights with mean one were assigned to patients and shared across all prediction points, index-age observations and generated futures belonging to the same individual. Level 1 retained its within-patient prediction-point weights, while generated futures retained a weight of 1*/M* within each index case. For Levels 1 and 2, the original panel of concepts with both outcome classes observed was held fixed when recalculating macro-AUROC and macro-AP, avoiding changes in the evaluated concept panel across replicates.

Uncertainty intervals were obtained from the 2.5th and 97.5th percentiles of the resulting Bayesian-bootstrap distributions. These intervals characterize uncertainty conditional on the evaluated patients, fixed concept panel, fitted models and saved predictions or generated futures. In Level 3, the same patient-level weighting scheme was applied when recomputing integrated survival, cumulative-incidence and transition-divergence measures.

To provide an empirical reference for interpreting trajectory-level discrepancies, observed patients were additionally partitioned into two disjoint groups in up to 100 repeated random splits per index-age cohort (seed 42 + age). The corresponding population-level distances were calculated between the two observed groups using the same metric definitions as in the observed–generated comparisons. The median and 2.5th–97.5th percentile range across valid partitions are reported together with the number of valid splits. These observed–observed comparisons characterize finite-sample variability in the recorded cohort and should not be interpreted as confidence intervals or universal lower bounds on generative model error.

### Dimension III: Representation Transferability

#### Level 4: Few-shot adaptation with a frozen pretrained backbone

Each task is performed using *K* labeled positive and *K* labeled negative examples, resulting in a total of 2*K* examples per draw. The protocol evaluates *K ∈ {*8, 16, 32, 64, 128*}* with 20 draws per setting. Developers extract the prescribed representations using their own model implementation, keep the pretrained backbone frozen and in evaluation mode, and fit a separate linear head for each support set. The repository supplies a reference head implementation. Cases without an observed target event within the prediction horizon are censored and excluded from both adaptation and test evaluation if follow-up is incomplete or unknown, or if death occurs within the horizon. For mortality tasks, death within the horizon is the target event and counts as a positive outcome. Tasks or settings with insufficient class support are retained as not estimable. The complete distribution of test performance across draws is retained, and summary statistics and confidence intervals are reported.

Each downstream task defines its own eligibility criteria, prediction index, clinical outcome, prediction horizon, and permitted event attributes. Numerical laboratory values may therefore be used when they form part of the clinical definition of a downstream task, even though laboratory values are not represented as separate concepts in the common autoregressive benchmark vocabulary. Tasks are bundled in catalogues that follow the template shown in Supplementary Table S6. In every catalogue, each task specifies anchor events, outcomes, and horizons. Laboratory mappings, measurement selection, and unit conversions remain developer responsibilities and must be declared. Level 4 tasks can be extended by creating new catalogues following the template.

For binary clinical outcomes, performance is summarized using AUROC and AP across *K*-shot conditions. Level 4 reports performance as a function of *K*, thereby characterizing the data efficiency with which pretrained representations can be adapted to new clinical prediction tasks.

The initial released catalogue contains 12 Type 2 Diabetes tasks (see Supplementary Table S5). Diagnosis anchors use the first recorded ICD-10 E11-family code. Selected code prefixes include the parent code and all descendants. For initiation tasks, patients with a known target before the anchor are excluded. Medication dispensing is preferred, with prescription accepted when dispensing is unavailable.

## Code availability

Reference code, the versioned benchmark clinical concept space, and evaluation implementations are available at https://dev.b2s.club/joana/ehrmeneia-benchmark. After local evaluation, EHRMENEIA automatically assembles the standardized aggregate results, metadata and support information into a portable benchmark package for visualization and submission, without including patient-level records, predictions, representations or generated trajectories.

The EHRMENEIA website https://ehrmeneia.upc.edu/ provides access to the benchmark results, model descriptions, graphical and numerical evaluation summaries, and downloadable information.

EHRMENEIA software is licensed under Apache-2.0. Original benchmark resources are licensed under CC BY 4.0, subject to third-party terminology restrictions.

## Data availability

The datasets used in this study were obtained through the official data releases and access procedures associated with the evaluated model implementations: EHRSHOT via Stanford data use agreement; MIMIC-IV via PhysioNet credentialed access; Delphi synthetic via the authors’ repository; Synthea OMOP example via the CEHR-GPT repository. No new patient-level clinical dataset was created or redistributed as part of EHRMENEIA. Access to each source dataset remains subject to the terms and procedures specified by its original provider. Aggregate EHRMENEIA evaluation results are made available through the benchmark website and associated repository.

## Acknowledgments

This work was supported by the Spanish Ministry of Economy and Competitiveness (MCIN/AEI/10.13039/501100011033) and European Union, project PID2025-173811OB-I00. We acknowledge the Networking Biomedical Research Centre in the subject area of Bioengineering, Biomaterials and Nanomedicine (CIBER-BBN), initiatives of Instituto de Investigación Carlos III (ISCIII). We acknowledge CERCA Programme/Generalitat de Catalunya. B2SLab is certified as 2025 SGR 00245. JF acknowledges the support from the Serra Húnter program. JGX acknowledges the support from the FPU Santander 2024 grant.

## Use of artificial intelligence

Generative AI tools were used to assist with code optimization and to improve the clarity and readability of the manuscript. All AI-assisted modifications were reviewed and validated by the authors, who take full responsibility for the final content.

