## Supplementary Materials for "EHRMENEIA: A Benchmark for Foundation Models on Longitudinal Electronic Health Records"

### S1 Model-specific implementation details

**CLMBR-T.** We evaluated the released CLMBR-T-base checkpoint using EHRSHOT data. The source comprised 6,739 patients, with 2,295 patients in the released training partition, 2,232 in tuning and 2,212 in held-out testing. The released training partition was excluded from evaluation; tuning patients were used for Level 4 adaptation and held-out patients for Level 4 testing and Level 1. The authors report pretraining on 2.57 million Stanford EHRs. The evaluated model contained 141,650,432 parameters, 12 Transformer layers and hidden dimension 768, with a 65,536-token input vocabulary, an 8,192-class next-token head and a maximum representation context of 496 native tokens. Level 1 sampling yielded 39,440 prediction points from 2,153 held-out patients. CLMBR-T exposed next-event predictions and pretrained representations but not a validated joint clinical-event and event-time generation interface, and was therefore evaluated at Levels 1 and 4 but not Levels 2–3.

**ETHOS.** We evaluated the original ETHOS checkpoint and its 4,367-token vocabulary using MIMIC-IV 2.2. The evaluated model contained 47,441,664 parameters, 6 Transformer layers, 12 attention heads, hidden dimension 768 and context length 2,048. The reconstructed held-out population contained 29,971 patients and was divided into 14,985 adaptation and 14,986 test patients. Level 1 sampling yielded 160,925 prediction points from 11,913 test patients. For Levels 2–3, the reference-age cohorts contained 288, 370 and 263 cases at ages 30, 50 and 80, respectively (921 longitudinal index cases total). Because MIMIC-IV does not provide complete dates of birth, they were approximated as 1 January of the estimated birth year; age-capped patients were excluded from longitudinal evaluations. We retained 100 generated futures per case. Of 92,100 retained generated futures, 6,129 contained a model-generated death event, 6,786 reached five years and 79,185 reached the native timeline-end state. Timeline end was treated as an absorbing no-further-record state and was not interpreted as death. The retained trajectories contained 16,132,370 mapped generated events.

**Delphi synthetic demonstration.** We evaluated the official Delphi demonstration architecture using the authors' distributed synthetic training and validation data. This was a locally trained synthetic implementation demonstration and not an evaluation of the original clinical Delphi checkpoint. The demonstration model contained 2,243,400 parameters, 12 layers, 12 attention heads, hidden dimension 120, context length 96 and 1,270 native output classes. Its benchmark projection contained diagnosis concepts only. Level 1 sampling yielded 52,378 prediction points from 2,858 patients. For Levels 2–3, the reference-age cohorts contained 1,110 and 2,216 cases at ages 30 and 50, respectively, with no eligible age-80 cases (3,326 longitudinal index cases total). We retained 100 generated futures per case, giving 332,600 trajectories; all retained trajectories reached five years or death under the native generator.

**CEHR-GPT synthetic demonstration.** We used the official CEHR-GPT implementation and the authors' Synthea OMOP example containing 1,141,631 synthetic patients. A fixed 42,000-patient subset was partitioned into 10,000 training, 2,000 tuning, 10,000 adaptation and 20,000 test patients; native preprocessing retained 9,987, 1,998, 9,984 and 19,970 patient sequences, respectively. The locally trained demonstration model contained 87,046,658 parameters, 12 GPT-2 blocks, 12 attention heads, hidden dimension 768, representation context 1,024 and a 1,796-token native vocabulary. Forecasting considered all 19,970 available native test sequences. Level 1 sampling yielded 106,294 prediction points from 16,174 patients. For Levels 2–3, this population yielded 9,772, 6,525 and 972 eligible longitudinal cases at ages 30, 50 and 80, respectively (17,269 cases from 10,925 distinct patients). One hundred futures per case yielded 1,726,900 retained trajectories; 1,409,118 reached five years and 317,782 terminated at native EOS. EOS was treated as an absorbing no-further-record state and not as death. The source contained no death table and no laboratory table; mortality and laboratory-dependent evaluations were therefore unsupported.

**Shared frozen-head fitting** All four applications used the same reference linear classifier fitted on representations extracted from a frozen backbone in evaluation mode. The pretrained backbone was kept fixed in evaluation mode, and a separate head was fitted for each support draw. Head weights and bias were initialized to zero. Features were normalized using the support-set population mean and standard deviation, with zero standard deviations replaced by one. The head was optimized by full-batch gradient descent for 2,000 updates with learning rate 0.01 and L2 coefficient 0.001.

Support sets contained $K$ positive and $K$ negative patients for $K \in \{8, 16, 32, 64, 128\}$. Twenty support draws were generated. Test evaluation required at least 20 positive and 20 negative cases; unsupported task–$K$ settings remained not estimable. AUROC and average precision were calculated on the fixed task-specific test cohort for each draw and summarized by the mean across draws. The released 95% confidence intervals used 2,000 percentile-bootstrap resamples of the 20 draw-level scores.

Table S1: **Evaluation scope across implementations.** In Level 1 prediction points are not distinct patients. Levels 2–3 use $M = 100$ futures per temporal case; cases are patient–index-age observations. Yes denotes an executed level. Level 4 counts tasks with at least one estimable $K$ and all estimable task–$K$ combinations, $K \in \{8, 16, 32, 64, 128\}$.

| Implementation | Evaluation setting | L1 | L2 | L3 | L4 | L1 patients | L1 points | Futures | L4 tasks / settings |
|---|---|---|---|---|---|---|---|---|---|
| CLMBR-T | Released clinical checkpoint; EHRSHOT | Yes | – | – | Yes | 2,153 | 39,440 | – | 10/12; 39 |
| ETHOS | Released clinical checkpoint; MIMIC-IV 2.2 | Yes | Yes | Yes | Yes | 11,913 | 160,925 | 92,100 | 10/12; 47 |
| Delphi demo | Official architecture; author-distributed synthetic data | Yes | Yes | Yes | Yes | 2,858 | 52,378 | 332,600 | 3/12; 9 |
| CEHR-GPT demo | Official architecture; Synthea OMOP example | Yes | Yes | Yes | Yes | 16,174 | 106,294 | 1,726,900 | 5/12; 21 |
| Total | – | – | – | – | – | 33,098 | 359,037 | 2,151,600 | 116 settings |

Table S2: **Level 1 next-event prediction.** Macro-AUROC and macro-AP are means across all computable concepts. Patient and prediction-point counts are distinct. To compute history, the reference, each benchmark concept was scored by its relative frequency among all clinical events retained in the model's native input before the prediction point. Parentheses give 95% patient-level Bayesian-bootstrap intervals (1,000 replicates), conditional on the original computable-concept panel and saved predictions. AP denotes average precision.

| Implementation | Patients | Points | Concepts | Macro-AUROC | Macro-AP | Top-5 | Top-10 |
|---|---|---|---|---|---|---|---|
| CEHR-GPT - synthetic demo | 16,174 | 106,294 | 89 | 0.5092 (0.5054–0.5144) | 0.0127 (0.0127–0.0130) | 0.247 | 0.641 |
| CEHR-GPT - synthetic demo / History | 16,174 | 106,294 | 89 | 0.5484 (0.5456–0.5517) | 0.0553 (0.0536–0.0582) | 0.581 | 0.583 |
| CLMBR-T-Base | 2,153 | 39,440 | 1,400 | 0.7058 (0.7037–0.7070) | 0.0612 (0.0601–0.0658) | 0.671 | 0.743 |
| CLMBR-T-Base / History | 2,153 | 39,440 | 1,400 | 0.7445 (0.7395–0.7453) | 0.1048 (0.1039–0.1093) | 0.500 | 0.616 |
| Delphi - synthetic demo | 2,858 | 52,378 | 829 | 0.6870 (0.6829–0.6916) | 0.0064 (0.0064–0.0081) | 0.138 | 0.223 |
| Delphi - synthetic demo / History | 2,858 | 52,378 | 829 | 0.4931 (0.4930–0.4932) | 0.0012 (0.0012–0.0012) | 0.000 | 0.000 |
| ETHOS | 11,913 | 160,925 | 1,697 | 0.5326 (0.5318–0.5332) | 0.0010 (0.0009–0.0013) | 0.370 | 0.450 |
| ETHOS / History | 11,913 | 160,925 | 1,697 | 0.5363 (0.5349–0.5378) | 0.0047 (0.0043–0.0056) | 0.067 | 0.151 |

Table S3: **Level 2 future-window prediction.** Brier is the unweighted mean of per-concept Brier scores. Marginal denotes prevalence from a separate patient cohort. Parentheses give 95% patient-level Bayesian-bootstrap intervals (1,000 replicates), conditional on the original computable-concept panel and saved predictions. AP denotes average precision.

| Implementation | Age | Horizon (d) | Cases | Concepts | Macro-AUROC | Macro-AP | Macro-Brier |
|---|---|---|---|---|---|---|---|
| CEHR-GPT - synthetic demo | 30 | 30 | 9,772 | 19 | 0.5644 (0.5390–0.5873) | 0.0220 (0.0047–0.0527) | 0.0003 |
| CEHR-GPT - synthetic demo / Marginal | 30 | 30 | 9,772 | 19 | 0.5000 (0.5000–0.5000) | 0.0003 (0.0002–0.0004) | 0.0003 |
| CEHR-GPT - synthetic demo | 30 | 90 | 9,772 | 35 | 0.5750 (0.5553–0.5951) | 0.0149 (0.0103–0.0249) | 0.0006 |
| CEHR-GPT - synthetic demo / Marginal | 30 | 90 | 9,772 | 35 | 0.5000 (0.5000–0.5000) | 0.0006 (0.0005–0.0007) | 0.0006 |
| CEHR-GPT - synthetic demo | 30 | 180 | 9,772 | 44 | 0.5506 (0.5362–0.5647) | 0.0424 (0.0392–0.0470) | 0.0026 |
| CEHR-GPT - synthetic demo / Marginal | 30 | 180 | 9,772 | 44 | 0.5000 (0.5000–0.5000) | 0.0029 (0.0028–0.0031) | 0.0028 |
| CEHR-GPT - synthetic demo | 30 | 365 | 9,772 | 51 | 0.5622 (0.5489–0.5764) | 0.0425 (0.0410–0.0458) | 0.0028 |
| CEHR-GPT - synthetic demo / Marginal | 30 | 365 | 9,772 | 51 | 0.5000 (0.5000–0.5000) | 0.0032 (0.0030–0.0034) | 0.0030 |
| CEHR-GPT - synthetic demo | 50 | 30 | 6,525 | 40 | 0.5602 (0.5481–0.5731) | 0.0679 (0.0557–0.0873) | 0.0028 |
| CEHR-GPT - synthetic demo / Marginal | 50 | 30 | 6,525 | 40 | 0.5000 (0.5000–0.5000) | 0.0029 (0.0027–0.0031) | 0.0028 |
| CEHR-GPT - synthetic demo | 50 | 90 | 6,525 | 57 | 0.5642 (0.5552–0.5736) | 0.0624 (0.0548–0.0721) | 0.0026 |
| CEHR-GPT - synthetic demo / Marginal | 50 | 90 | 6,525 | 57 | 0.5000 (0.5000–0.5000) | 0.0029 (0.0027–0.0031) | 0.0027 |
| CEHR-GPT - synthetic demo | 50 | 180 | 6,525 | 59 | 0.5589 (0.5493–0.5687) | 0.0620 (0.0541–0.0714) | 0.0030 |
| CEHR-GPT - synthetic demo / Marginal | 50 | 180 | 6,525 | 59 | 0.5000 (0.5000–0.5000) | 0.0034 (0.0032–0.0036) | 0.0032 |
| CEHR-GPT - synthetic demo | 50 | 365 | 6,525 | 74 | 0.5597 (0.5498–0.5701) | 0.0673 (0.0625–0.0737) | 0.0040 |
| CEHR-GPT - synthetic demo / Marginal | 50 | 365 | 6,525 | 74 | 0.5000 (0.5000–0.5000) | 0.0049 (0.0046–0.0051) | 0.0044 |
| CEHR-GPT - synthetic demo | 80 | 30 | 972 | 15 | 0.6735 (0.6638–0.6802) | 0.2201 (0.1766–0.2688) | 0.0052 |
| CEHR-GPT - synthetic demo / Marginal | 80 | 30 | 972 | 15 | 0.5000 (0.5000–0.5000) | 0.0069 (0.0052–0.0087) | 0.0068 |
| CEHR-GPT - synthetic demo | 80 | 90 | 972 | 28 | 0.6010 (0.5829–0.6194) | 0.1410 (0.1297–0.1578) | 0.0115 |
| CEHR-GPT - synthetic demo / Marginal | 80 | 90 | 972 | 28 | 0.5000 (0.5000–0.5000) | 0.0154 (0.0134–0.0174) | 0.0143 |
| CEHR-GPT - synthetic demo | 80 | 180 | 972 | 42 | 0.5956 (0.5819–0.6086) | 0.1323 (0.1212–0.1429) | 0.0147 |
| CEHR-GPT - synthetic demo / Marginal | 80 | 180 | 972 | 42 | 0.5000 (0.5000–0.5000) | 0.0220 (0.0201–0.0240) | 0.0184 |
| CEHR-GPT - synthetic demo | 80 | 365 | 972 | 46 | 0.6026 (0.5889–0.6167) | 0.1260 (0.1155–0.1355) | 0.0130 |
| CEHR-GPT - synthetic demo / Marginal | 80 | 365 | 972 | 46 | 0.5000 (0.5000–0.5000) | 0.0230 (0.0210–0.0250) | 0.0190 |
| Delphi - synthetic demo | 30 | 30 | 1,110 | 7 | 0.4981 (0.4973–0.4987) | 0.0009 (0.0004–0.0017) | 0.0009 |
| Delphi - synthetic demo / Marginal | 30 | 30 | 1,110 | 7 | 0.5000 (0.5000–0.5000) | 0.0009 (0.0004–0.0017) | 0.0009 |
| Delphi - synthetic demo | 30 | 90 | 1,110 | 14 | 0.4929 (0.4918–0.4938) | 0.0010 (0.0006–0.0016) | 0.0010 |
| Delphi - synthetic demo / Marginal | 30 | 90 | 1,110 | 14 | 0.5000 (0.5000–0.5000) | 0.0010 (0.0006–0.0016) | 0.0010 |
| Delphi - synthetic demo | 30 | 180 | 1,110 | 28 | 0.5131 (0.5073–0.5219) | 0.0030 (0.0011–0.0069) | 0.0011 |
| Delphi - synthetic demo / Marginal | 30 | 180 | 1,110 | 28 | 0.5000 (0.5000–0.5000) | 0.0011 (0.0008–0.0016) | 0.0011 |
| Delphi - synthetic demo | 30 | 365 | 1,110 | 60 | 0.5278 (0.5196–0.5353) | 0.0061 (0.0029–0.0118) | 0.0012 |
| Delphi - synthetic demo / Marginal | 30 | 365 | 1,110 | 60 | 0.5000 (0.5000–0.5000) | 0.0012 (0.0009–0.0015) | 0.0012 |
| Delphi - synthetic demo | 50 | 30 | 2,216 | 52 | 0.5185 (0.5126–0.5250) | 0.0020 (0.0008–0.0049) | 0.0005 |
| Delphi - synthetic demo / Marginal | 50 | 30 | 2,216 | 52 | 0.5000 (0.5000–0.5000) | 0.0005 (0.0004–0.0007) | 0.0005 |
| Delphi - synthetic demo | 50 | 90 | 2,216 | 109 | 0.5216 (0.5148–0.5287) | 0.0016 (0.0011–0.0025) | 0.0007 |
| Delphi - synthetic demo / Marginal | 50 | 90 | 2,216 | 109 | 0.5000 (0.5000–0.5000) | 0.0007 (0.0006–0.0009) | 0.0007 |
| Delphi - synthetic demo | 50 | 180 | 2,216 | 154 | 0.5187 (0.5120–0.5257) | 0.0020 (0.0016–0.0036) | 0.0010 |
| Delphi - synthetic demo / Marginal | 50 | 180 | 2,216 | 154 | 0.5000 (0.5000–0.5000) | 0.0010 (0.0009–0.0011) | 0.0010 |
| Delphi - synthetic demo | 50 | 365 | 2,216 | 233 | 0.5325 (0.5250–0.5399) | 0.0061 (0.0045–0.0106) | 0.0013 |
| Delphi - synthetic demo / Marginal | 50 | 365 | 2,216 | 233 | 0.5000 (0.5000–0.5000) | 0.0013 (0.0012–0.0014) | 0.0013 |
| Delphi - synthetic demo | 80 | 30 | 0 | 0 | NA | NA | NA |
| Delphi - synthetic demo / Marginal | 80 | 30 | 0 | 0 | NA | NA | NA |
| Delphi - synthetic demo | 80 | 90 | 0 | 0 | NA | NA | NA |
| Delphi - synthetic demo / Marginal | 80 | 90 | 0 | 0 | NA | NA | NA |
| Delphi - synthetic demo | 80 | 180 | 0 | 0 | NA | NA | NA |
| Delphi - synthetic demo / Marginal | 80 | 180 | 0 | 0 | NA | NA | NA |
| Delphi - synthetic demo | 80 | 365 | 0 | 0 | NA | NA | NA |
| Delphi - synthetic demo / Marginal | 80 | 365 | 0 | 0 | NA | NA | NA |
| ETHOS | 30 | 30 | 288 | 130 | 0.6216 (0.5962–0.6456) | 0.0960 (0.0723–0.1334) | 0.0210 |
| ETHOS / Marginal | 30 | 30 | 288 | 130 | 0.5000 (0.5000–0.5000) | 0.0180 (0.0121–0.0244) | 0.0169 |
| ETHOS | 30 | 90 | 288 | 256 | 0.6572 (0.6428–0.6718) | 0.1430 (0.1211–0.1748) | 0.0219 |
| ETHOS / Marginal | 30 | 90 | 288 | 256 | 0.5000 (0.5000–0.5000) | 0.0237 (0.0186–0.0294) | 0.0203 |
| ETHOS | 30 | 180 | 288 | 313 | 0.6913 (0.6754–0.7076) | 0.1853 (0.1632–0.2246) | 0.0254 |
| ETHOS / Marginal | 30 | 180 | 288 | 313 | 0.5000 (0.5000–0.5000) | 0.0297 (0.0245–0.0358) | 0.0244 |
| ETHOS | 30 | 365 | 288 | 435 | 0.6877 (0.6714–0.7036) | 0.2064 (0.1833–0.2536) | 0.0289 |
| ETHOS / Marginal | 30 | 365 | 288 | 435 | 0.5000 (0.5000–0.5000) | 0.0368 (0.0316–0.0425) | 0.0266 |
| ETHOS | 50 | 30 | 370 | 282 | 0.7611 (0.7477–0.7728) | 0.2443 (0.2161–0.2868) | 0.0154 |
| ETHOS / Marginal | 50 | 30 | 370 | 282 | 0.5000 (0.5000–0.5000) | 0.0151 (0.0113–0.0197) | 0.0138 |
| ETHOS | 50 | 90 | 370 | 470 | 0.7286 (0.7150–0.7415) | 0.2291 (0.2085–0.2653) | 0.0165 |
| ETHOS / Marginal | 50 | 90 | 370 | 470 | 0.5000 (0.5000–0.5000) | 0.0193 (0.0158–0.0236) | 0.0162 |
| ETHOS | 50 | 180 | 370 | 669 | 0.7117 (0.7000–0.7230) | 0.2252 (0.2051–0.2608) | 0.0179 |
| ETHOS / Marginal | 50 | 180 | 370 | 669 | 0.5000 (0.5000–0.5000) | 0.0226 (0.0192–0.0269) | 0.0173 |
| ETHOS | 50 | 365 | 370 | 827 | 0.7113 (0.6985–0.7244) | 0.2345 (0.2159–0.2681) | 0.0212 |
| ETHOS / Marginal | 50 | 365 | 370 | 827 | 0.5000 (0.5000–0.5000) | 0.0291 (0.0255–0.0334) | 0.0196 |

| Implementation | Age | Horizon (d) | Cases | Concepts | Macro-AUROC | Macro-AP | Macro-Brier |
|---|---|---|---|---|---|---|---|
| ETHOS | 80 | 30 | 263 | 250 | 0.7409 (0.7212–0.7595) | 0.1562 (0.1164–0.2137) | 0.0215 |
| ETHOS / Marginal | 80 | 30 | 263 | 250 | 0.5000 (0.5000–0.5000) | 0.0166 (0.0114–0.0231) | 0.0155 |
| ETHOS | 80 | 90 | 263 | 401 | 0.6904 (0.6706–0.7093) | 0.1603 (0.1364–0.2001) | 0.0258 |
| ETHOS / Marginal | 80 | 90 | 263 | 401 | 0.5000 (0.5000–0.5000) | 0.0285 (0.0228–0.0345) | 0.0227 |
| ETHOS | 80 | 180 | 263 | 652 | 0.6711 (0.6570–0.6843) | 0.1747 (0.1615–0.2018) | 0.0281 |
| ETHOS / Marginal | 80 | 180 | 263 | 652 | 0.5000 (0.5000–0.5000) | 0.0355 (0.0302–0.0414) | 0.0255 |
| ETHOS | 80 | 365 | 263 | 823 | 0.6582 (0.6447–0.6718) | 0.1966 (0.1861–0.2219) | 0.0324 |
| ETHOS / Marginal | 80 | 365 | 263 | 823 | 0.5000 (0.5000–0.5000) | 0.0437 (0.0381–0.0493) | 0.0290 |

Table S4: **Level 3 population-level fidelity and observed–observed references.** Distances quantify discrepancies between observed and generated trajectories in survival, cumulative incidence, and sequential event transitions. Parentheses accompanying these distances indicate 95% patient-level Bayesian-bootstrap uncertainty intervals, conditional on the evaluated cohort and generated futures. Observed–observed values summarize repeated split-half comparisons of the observed cohort, with parentheses indicating the 2.5th–97.5th percentile range across valid partitions. Survival and competing-risk incidence estimates require appropriate mortality observations.

| Implementation | Age | Component | Cases | Futures | Distance (95% interval) | Observed–observed | Valid splits |
|---|---|---|---|---|---|---|---|
| CEHR-GPT - synthetic demo | 30 | survival | 9,772 | 977,200 | 0.0000 (0.0000–0.0000) | 0.0000 (0.0000–0.0000) | 100 |
| CEHR-GPT - synthetic demo | 30 | incidence E11 | 9,759 | 975,900 | 0.0004 ($8.77 \times 10^{-5}$–0.0008) | 0.0003 (0.0001–0.0010) | 100 |
| CEHR-GPT - synthetic demo | 30 | incidence J44 | 9,772 | 977,200 | 0.0000 (0.0000–0.0000) | 0.0000 (0.0000–0.0000) | 100 |
| CEHR-GPT - synthetic demo | 30 | incidence C34 | 9,772 | 977,200 | 0.0000 (0.0000–0.0000) | 0.0000 (0.0000–0.0000) | 100 |
| CEHR-GPT - synthetic demo | 30 | incidence F32 | 9,772 | 977,200 | 0.0000 (0.0000–0.0000) | 0.0000 (0.0000–0.0000) | 100 |
| CEHR-GPT - synthetic demo | 30 | incidence N17 | 9,772 | 977,200 | 0.0000 (0.0000–0.0000) | 0.0000 (0.0000–0.0000) | 100 |
| CEHR-GPT - synthetic demo | 30 | incidence I21 | 9,772 | 977,200 | 0.0000 (0.0000–0.0000) | 0.0000 (0.0000–0.0000) | 100 |
| CEHR-GPT - synthetic demo | 30 | transitions | 9,772 | 977,200 | 0.3256 (0.2671–0.3965) | 0.1836 (0.1080–0.2771) | 100 |
| CEHR-GPT - synthetic demo | 50 | survival | 6,525 | 652,500 | 0.0000 (0.0000–0.0000) | 0.0000 (0.0000–0.0000) | 100 |
| CEHR-GPT - synthetic demo | 50 | incidence E11 | 6,453 | 645,300 | 0.0020 (0.0010–0.0033) | 0.0008 (0.0003–0.0024) | 100 |
| CEHR-GPT - synthetic demo | 50 | incidence J44 | 6,525 | 652,500 | 0.0000 (0.0000–0.0000) | 0.0000 (0.0000–0.0000) | 100 |
| CEHR-GPT - synthetic demo | 50 | incidence C34 | 6,525 | 652,500 | 0.0000 (0.0000–0.0000) | 0.0000 (0.0000–0.0000) | 100 |
| CEHR-GPT - synthetic demo | 50 | incidence F32 | 6,525 | 652,500 | 0.0000 (0.0000–0.0000) | 0.0000 (0.0000–0.0000) | 100 |
| CEHR-GPT - synthetic demo | 50 | incidence N17 | 6,525 | 652,500 | 0.0000 (0.0000–0.0000) | 0.0000 (0.0000–0.0000) | 100 |
| CEHR-GPT - synthetic demo | 50 | incidence I21 | 6,525 | 652,500 | 0.0000 (0.0000–0.0000) | 0.0000 (0.0000–0.0000) | 100 |
| CEHR-GPT - synthetic demo | 50 | transitions | 6,525 | 652,500 | 0.3763 (0.3221–0.4534) | 0.0967 (0.0579–0.2488) | 100 |
| CEHR-GPT - synthetic demo | 80 | survival | 972 | 97,200 | 0.0000 (0.0000–0.0000) | 0.0000 (0.0000–0.0000) | 100 |
| CEHR-GPT - synthetic demo | 80 | incidence E11 | 939 | 93,900 | 0.0012 (0.0010–0.0014) | $1.55 \times 10^{-5}$ ($1.47 \times 10^{-5}$–$1.63 \times 10^{-5}$) | 100 |
| CEHR-GPT - synthetic demo | 80 | incidence J44 | 972 | 97,200 | 0.0000 (0.0000–0.0000) | 0.0000 (0.0000–0.0000) | 100 |
| CEHR-GPT - synthetic demo | 80 | incidence C34 | 972 | 97,200 | 0.0000 (0.0000–0.0000) | 0.0000 (0.0000–0.0000) | 100 |
| CEHR-GPT - synthetic demo | 80 | incidence F32 | 972 | 97,200 | 0.0000 (0.0000–0.0000) | 0.0000 (0.0000–0.0000) | 100 |
| CEHR-GPT - synthetic demo | 80 | incidence N17 | 972 | 97,200 | 0.0000 (0.0000–0.0000) | 0.0000 (0.0000–0.0000) | 100 |
| CEHR-GPT - synthetic demo | 80 | incidence I21 | 972 | 97,200 | 0.0000 (0.0000–0.0000) | 0.0000 (0.0000–0.0000) | 100 |
| CEHR-GPT - synthetic demo | 80 | transitions | 972 | 97,200 | 0.4043 (0.3446–0.6596) | 1.0000 (1.0000–1.0000) | 59 |
| Delphi - synthetic demo | 30 | survival | 1,110 | 111,000 | $1.34 \times 10^{-6}$ ($3.32 \times 10^{-8}$–$5.15 \times 10^{-6}$) | 0.0000 (0.0000–0.0000) | 100 |
| Delphi - synthetic demo | 30 | incidence E11 | 1,110 | 111,000 | $5.98 \times 10^{-5}$ ($3.09 \times 10^{-5}$–$9.35 \times 10^{-5}$) | 0.0000 (0.0000–0.0000) | 100 |
| Delphi - synthetic demo | 30 | incidence J44 | 1,110 | 111,000 | $2.66 \times 10^{-5}$ ($7.95 \times 10^{-6}$–$5.81 \times 10^{-5}$) | 0.0000 (0.0000–0.0000) | 100 |
| Delphi - synthetic demo | 30 | incidence C34 | 1,110 | 111,000 | 0.0000 (0.0000–0.0000) | 0.0000 (0.0000–0.0000) | 100 |
| Delphi - synthetic demo | 30 | incidence F32 | 1,062 | 106,200 | 0.0008 (0.0006–0.0042) | 0.0021 (0.0005–0.0065) | 100 |
| Delphi - synthetic demo | 30 | incidence N17 | 1,110 | 111,000 | $1.01 \times 10^{-5}$ ($1.10 \times 10^{-6}$–$2.95 \times 10^{-5}$) | 0.0000 (0.0000–0.0000) | 100 |

| Implementation | Age | Component | Cases | Futures | Distance (95% interval) | Observed–observed | Valid splits |
|---|---|---|---|---|---|---|---|
| Delphi - synthetic demo | 30 | incidence I21 | 1,109 | 110,900 | 0.0008 (0.0007–0.0009) | 0.0000 (0.0000–0.0000) | 100 |
| Delphi - synthetic demo | 30 | transitions | 1,110 | 111,000 | 0.9572 (0.9198–0.9913) | 1.0000 (1.0000–1.0000) | 82 |
| Delphi - synthetic demo | 50 | survival | 2,216 | 221,600 | 0.0002 (0.0001–0.0004) | $4.99 \times 10^{-5}$ ($4.99 \times 10^{-5}$–0.0002) | 100 |
| Delphi - synthetic demo | 50 | incidence E11 | 2,194 | 219,400 | 0.0023 (0.0006–0.0052) | 0.0021 (0.0007–0.0054) | 100 |
| Delphi - synthetic demo | 50 | incidence J44 | 2,214 | 221,400 | 0.0006 (0.0003–0.0012) | 0.0009 (0.0004–0.0018) | 100 |
| Delphi - synthetic demo | 50 | incidence C34 | 2,214 | 221,400 | 0.0001 ($3.56 \times 10^{-5}$–0.0004) | 0.0002 (0.0002–0.0002) | 100 |
| Delphi - synthetic demo | 50 | incidence F32 | 2,040 | 204,000 | 0.0017 (0.0006–0.0055) | 0.0027 (0.0012–0.0080) | 100 |
| Delphi - synthetic demo | 50 | incidence N17 | 2,215 | 221,500 | 0.0002 ($4.91 \times 10^{-5}$–0.0010) | 0.0006 (0.0006–0.0006) | 100 |
| Delphi - synthetic demo | 50 | incidence I21 | 2,196 | 219,600 | 0.0017 (0.0007–0.0031) | 0.0016 (0.0006–0.0038) | 100 |
| Delphi - synthetic demo | 50 | transitions | 2,216 | 221,600 | 0.7266 (0.7156–0.8043) | 0.8822 (0.8013–0.9608) | 100 |
| Delphi - synthetic demo | 80 | survival (not computable) | 0 | 0 | NA | NA | 0 |
| Delphi - synthetic demo | 80 | incidence E11 (not computable) | 0 | 0 | NA | NA | 0 |
| Delphi - synthetic demo | 80 | incidence J44 (not computable) | 0 | 0 | NA | NA | 0 |
| Delphi - synthetic demo | 80 | incidence C34 (not computable) | 0 | 0 | NA | NA | 0 |
| Delphi - synthetic demo | 80 | incidence F32 (not computable) | 0 | 0 | NA | NA | 0 |
| Delphi - synthetic demo | 80 | incidence N17 (not computable) | 0 | 0 | NA | NA | 0 |
| Delphi - synthetic demo | 80 | incidence I21 (not computable) | 0 | 0 | NA | NA | 0 |
| Delphi - synthetic demo | 80 | transitions (not computable) | 0 | 0 | NA | NA | 0 |
| ETHOS | 30 | survival | 288 | 28,800 | 0.0015 (0.0010–0.0092) | 0.0066 (0.0060–0.0072) | 100 |
| ETHOS | 30 | incidence E11 | 283 | 28,300 | 0.0046 (0.0017–0.0164) | 0.0085 (0.0062–0.0183) | 100 |
| ETHOS | 30 | incidence J44 | 287 | 28,700 | 0.0029 (0.0009–0.0100) | 0.0058 (0.0052–0.0066) | 100 |
| ETHOS | 30 | incidence C34 | 288 | 28,800 | 0.0004 (0.0002–0.0006) | 0.0000 (0.0000–0.0000) | 100 |
| ETHOS | 30 | incidence F32 | 266 | 26,600 | 0.0224 (0.0114–0.0447) | 0.0227 (0.0097–0.0549) | 100 |
| ETHOS | 30 | incidence N17 | 286 | 28,600 | 0.0048 (0.0027–0.0143) | 0.0096 (0.0049–0.0118) | 100 |
| ETHOS | 30 | incidence I21 | 288 | 28,800 | 0.0010 (0.0005–0.0016) | 0.0000 (0.0000–0.0000) | 100 |
| ETHOS | 30 | transitions | 288 | 28,800 | 0.4922 (0.4489–0.5383) | 0.1641 (0.1027–0.3513) | 100 |
| ETHOS | 50 | survival | 370 | 37,000 | 0.0143 (0.0051–0.0334) | 0.0163 (0.0064–0.0430) | 100 |
| ETHOS | 50 | incidence E11 | 321 | 32,100 | 0.0083 (0.0046–0.0209) | 0.0171 (0.0062–0.0469) | 100 |
| ETHOS | 50 | incidence J44 | 364 | 36,400 | 0.0076 (0.0036–0.0162) | 0.0109 (0.0044–0.0243) | 100 |
| ETHOS | 50 | incidence C34 | 370 | 37,000 | 0.0019 (0.0006–0.0078) | 0.0043 (0.0039–0.0046) | 100 |

| Implementation | Age | Component | Cases | Futures | Distance (95% interval) | Observed–observed | Valid splits |
|---|---|---|---|---|---|---|---|
| ETHOS | 50 | incidence F32 | 321 | 32,100 | 0.0308 (0.0177–0.0582) | 0.0289 (0.0130–0.0759) | 100 |
| ETHOS | 50 | incidence N17 | 337 | 33,700 | 0.0130 (0.0047–0.0327) | 0.0174 (0.0081–0.0435) | 100 |
| ETHOS | 50 | incidence I21 | 363 | 36,300 | 0.0021 (0.0014–0.0132) | 0.0083 (0.0056–0.0201) | 100 |
| ETHOS | 50 | transitions | 370 | 37,000 | 0.4311 (0.4235–0.4789) | 0.1635 (0.1285–0.2187) | 100 |
| ETHOS | 80 | survival | 263 | 26,300 | 0.0515 (0.0297–0.0846) | 0.0284 (0.0116–0.0956) | 100 |
| ETHOS | 80 | incidence E11 | 180 | 18,000 | 0.0174 (0.0064–0.0573) | 0.0360 (0.0081–0.0918) | 100 |
| ETHOS | 80 | incidence J44 | 237 | 23,700 | 0.0212 (0.0079–0.0523) | 0.0221 (0.0049–0.0661) | 100 |
| ETHOS | 80 | incidence C34 | 257 | 25,700 | 0.0072 (0.0051–0.0126) | 0.0046 (0.0041–0.0053) | 100 |
| ETHOS | 80 | incidence F32 | 228 | 22,800 | 0.0304 (0.0128–0.0624) | 0.0300 (0.0115–0.0738) | 100 |
| ETHOS | 80 | incidence N17 | 212 | 21,200 | 0.0609 (0.0311–0.1048) | 0.0355 (0.0139–0.0961) | 100 |
| ETHOS | 80 | incidence I21 | 249 | 24,900 | 0.0245 (0.0077–0.0537) | 0.0205 (0.0087–0.0590) | 100 |
| ETHOS | 80 | transitions | 263 | 26,300 | 0.3812 (0.3742–0.4260) | 0.1409 (0.1134–0.1966) | 100 |

Table S5: **Released Level 4 type 2 diabetes catalogue.** Diagnosis and medication definitions use the benchmark code families shown; selected code prefixes include their eligible descendants.

| Task family | Anchor event | Target outcome | Horizon |
|---|---|---|---|
| Biomarker-to-Onset | HbA1c $\leq$ 6% | E11 diagnosis | 5 years |
| | 6% $<$ HbA1c $<$ 7.5% | E11 diagnosis | 5 years |
| | HbA1c $\geq$ 7.5% | E11 diagnosis | 2 years |
| Onset-to-Medication | E11 diagnosis | Non-insulin glucose-lowering medication (A10B) | 2 years |
| | E11 diagnosis | Insulin or analogue (A10A) | 2 years |
| | E11 diagnosis | Insulin or analogue (A10A) | 7 years |
| Onset-to-Comorbidity | E11 diagnosis | Chronic kidney disease diagnosis (N18) | 2 years |
| | E11 diagnosis | Chronic kidney disease diagnosis (N18) | 10 years |
| | E11 diagnosis | Retinal disorder diagnosis (H36) | 5 years |
| | E11 diagnosis | Neuropathy diagnosis (G63 or G62.9) | 5 years |
| | E11 diagnosis | Cardiovascular diagnosis composite (I25, I50, or I63) | 5 years |
| Onset-to-Death | E11 diagnosis | All-cause mortality | 10 years |

Table S6: **Level 4 catalogue template.** A Level 4 catalogue defines clinically interpretable task families through an anchor, a target outcome and a prediction horizon. The rows illustrate supported catalogue structures.

| **Task template** | **Anchor** | **Target outcome** |
|---|---|---|
| Biomarker-to-Onset | Laboratory measurement or biomarker state | Subsequent recorded disease diagnosis |
| Onset-to-Medication | Index disease diagnosis | Initiation of a relevant pharmacological treatment |
| Onset-to-Comorbidity | Index disease diagnosis | Subsequent related diagnosis or complication |
| Onset-to-Death | Index disease diagnosis | All-cause mortality |

Table S7: **Level 4 representation transfer across all task–$K$ settings.** Test counts include the uncensored test cases. Parentheses give 95% percentile-bootstrap intervals for the mean over 20 support draws, conditional on the fixed test cohort. All 12 tasks are reported for every implementation at $K \in \{8, 16, 32, 64, 128\}$, including non-estimable combinations.

| Implementation | Task | $K$ | Test $n$ | Positive $n$ | AUROC (95% CI) | AP (95% CI) |
|---|---|---|---|---|---|---|
| CLMBR-T | HbA1c ≤6% to E11, 5 y | 8 | 308 | 87 | 0.695 (0.674–0.713) | 0.492 (0.463–0.517) |
| CLMBR-T | HbA1c ≤6% to E11, 5 y | 16 | 308 | 87 | 0.711 (0.692–0.728) | 0.500 (0.471–0.526) |
| CLMBR-T | HbA1c ≤6% to E11, 5 y | 32 | 308 | 87 | 0.738 (0.724–0.753) | 0.569 (0.548–0.591) |
| CLMBR-T | HbA1c ≤6% to E11, 5 y | 64 | 308 | 87 | 0.787 (0.780–0.796) | 0.640 (0.629–0.651) |
| CLMBR-T | HbA1c ≤6% to E11, 5 y | 128 | 308 | 87 | Not estimable | Not estimable |
| CLMBR-T | HbA1c >6 to <7.5% to E11, 5 y | 8 | 140 | 90 | 0.773 (0.744–0.797) | 0.852 (0.826–0.874) |
| CLMBR-T | HbA1c >6 to <7.5% to E11, 5 y | 16 | 140 | 90 | 0.815 (0.792–0.836) | 0.889 (0.868–0.905) |
| CLMBR-T | HbA1c >6 to <7.5% to E11, 5 y | 32 | 140 | 90 | 0.843 (0.825–0.858) | 0.908 (0.897–0.918) |
| CLMBR-T | HbA1c >6 to <7.5% to E11, 5 y | 64 | 140 | 90 | Not estimable | Not estimable |
| CLMBR-T | HbA1c >6 to <7.5% to E11, 5 y | 128 | 140 | 90 | Not estimable | Not estimable |
| CLMBR-T | HbA1c ≥7.5% to E11, 2 y | 8 | 92 | 60 | 0.882 (0.867–0.896) | 0.902 (0.885–0.918) |
| CLMBR-T | HbA1c ≥7.5% to E11, 2 y | 16 | 92 | 60 | 0.875 (0.862–0.888) | 0.888 (0.871–0.904) |
| CLMBR-T | HbA1c ≥7.5% to E11, 2 y | 32 | 92 | 60 | Not estimable | Not estimable |
| CLMBR-T | HbA1c ≥7.5% to E11, 2 y | 64 | 92 | 60 | Not estimable | Not estimable |
| CLMBR-T | HbA1c ≥7.5% to E11, 2 y | 128 | 92 | 60 | Not estimable | Not estimable |
| CLMBR-T | E11 to A10B, 2 y | 8 | 256 | 100 | 0.522 (0.503–0.542) | 0.419 (0.403–0.436) |
| CLMBR-T | E11 to A10B, 2 y | 16 | 256 | 100 | 0.558 (0.538–0.579) | 0.440 (0.421–0.457) |
| CLMBR-T | E11 to A10B, 2 y | 32 | 256 | 100 | 0.581 (0.565–0.596) | 0.456 (0.441–0.470) |
| CLMBR-T | E11 to A10B, 2 y | 64 | 256 | 100 | 0.593 (0.581–0.605) | 0.452 (0.441–0.463) |
| CLMBR-T | E11 to A10B, 2 y | 128 | 256 | 100 | Not estimable | Not estimable |
| CLMBR-T | E11 to A10A, 2 y | 8 | 196 | 101 | 0.632 (0.605–0.657) | 0.649 (0.621–0.676) |
| CLMBR-T | E11 to A10A, 2 y | 16 | 196 | 101 | 0.659 (0.639–0.677) | 0.674 (0.650–0.698) |
| CLMBR-T | E11 to A10A, 2 y | 32 | 196 | 101 | 0.674 (0.655–0.692) | 0.694 (0.671–0.717) |
| CLMBR-T | E11 to A10A, 2 y | 64 | 196 | 101 | 0.710 (0.700–0.721) | 0.749 (0.736–0.761) |
| CLMBR-T | E11 to A10A, 2 y | 128 | 196 | 101 | Not estimable | Not estimable |
| CLMBR-T | E11 to A10A, 7 y | 8 | 173 | 135 | 0.696 (0.678–0.712) | 0.893 (0.884–0.901) |
| CLMBR-T | E11 to A10A, 7 y | 16 | 173 | 135 | 0.707 (0.687–0.724) | 0.895 (0.886–0.904) |
| CLMBR-T | E11 to A10A, 7 y | 32 | 173 | 135 | 0.722 (0.705–0.738) | 0.905 (0.897–0.914) |
| CLMBR-T | E11 to A10A, 7 y | 64 | 173 | 135 | Not estimable | Not estimable |
| CLMBR-T | E11 to A10A, 7 y | 128 | 173 | 135 | Not estimable | Not estimable |
| CLMBR-T | E11 to N18, 2 y | 8 | 303 | 75 | 0.676 (0.656–0.695) | 0.468 (0.432–0.499) |
| CLMBR-T | E11 to N18, 2 y | 16 | 303 | 75 | 0.698 (0.684–0.712) | 0.507 (0.488–0.525) |
| CLMBR-T | E11 to N18, 2 y | 32 | 303 | 75 | 0.729 (0.710–0.746) | 0.548 (0.519–0.573) |
| CLMBR-T | E11 to N18, 2 y | 64 | 303 | 75 | 0.757 (0.749–0.765) | 0.595 (0.581–0.608) |
| CLMBR-T | E11 to N18, 2 y | 128 | 303 | 75 | Not estimable | Not estimable |
| CLMBR-T | E11 to N18, 10 y | 8 | 162 | 123 | 0.693 (0.659–0.724) | 0.880 (0.862–0.896) |
| CLMBR-T | E11 to N18, 10 y | 16 | 162 | 123 | 0.747 (0.727–0.764) | 0.908 (0.899–0.916) |
| CLMBR-T | E11 to N18, 10 y | 32 | 162 | 123 | 0.774 (0.761–0.787) | 0.922 (0.916–0.928) |
| CLMBR-T | E11 to N18, 10 y | 64 | 162 | 123 | Not estimable | Not estimable |
| CLMBR-T | E11 to N18, 10 y | 128 | 162 | 123 | Not estimable | Not estimable |
| CLMBR-T | E11 to H36, 5 y | 8 | 178 | 0 | Not estimable | Not estimable |
| CLMBR-T | E11 to H36, 5 y | 16 | 178 | 0 | Not estimable | Not estimable |
| CLMBR-T | E11 to H36, 5 y | 32 | 178 | 0 | Not estimable | Not estimable |
| CLMBR-T | E11 to H36, 5 y | 64 | 178 | 0 | Not estimable | Not estimable |
| CLMBR-T | E11 to H36, 5 y | 128 | 178 | 0 | Not estimable | Not estimable |
| CLMBR-T | E11 to G63/G62.9, 5 y | 8 | 179 | 1 | Not estimable | Not estimable |
| CLMBR-T | E11 to G63/G62.9, 5 y | 16 | 179 | 1 | Not estimable | Not estimable |
| CLMBR-T | E11 to G63/G62.9, 5 y | 32 | 179 | 1 | Not estimable | Not estimable |
| CLMBR-T | E11 to G63/G62.9, 5 y | 64 | 179 | 1 | Not estimable | Not estimable |
| CLMBR-T | E11 to G63/G62.9, 5 y | 128 | 179 | 1 | Not estimable | Not estimable |
| CLMBR-T | E11 to I25/I50/I63, 5 y | 8 | 211 | 123 | 0.650 (0.633–0.666) | 0.727 (0.708–0.747) |
| CLMBR-T | E11 to I25/I50/I63, 5 y | 16 | 211 | 123 | 0.672 (0.650–0.690) | 0.751 (0.730–0.768) |
| CLMBR-T | E11 to I25/I50/I63, 5 y | 32 | 211 | 123 | 0.695 (0.684–0.707) | 0.774 (0.763–0.784) |
| CLMBR-T | E11 to I25/I50/I63, 5 y | 64 | 211 | 123 | 0.702 (0.693–0.710) | 0.780 (0.774–0.788) |
| CLMBR-T | E11 to I25/I50/I63, 5 y | 128 | 211 | 123 | Not estimable | Not estimable |
| CLMBR-T | E11 to death, 10 y | 8 | 144 | 81 | 0.754 (0.722–0.783) | 0.807 (0.779–0.832) |
| CLMBR-T | E11 to death, 10 y | 16 | 144 | 81 | 0.800 (0.775–0.822) | 0.848 (0.828–0.866) |
| CLMBR-T | E11 to death, 10 y | 32 | 144 | 81 | 0.803 (0.784–0.820) | 0.854 (0.840–0.868) |
| CLMBR-T | E11 to death, 10 y | 64 | 144 | 81 | 0.835 (0.831–0.841) | 0.886 (0.884–0.889) |
| CLMBR-T | E11 to death, 10 y | 128 | 144 | 81 | Not estimable | Not estimable |
| ETHOS | HbA1c ≤6% to E11, 5 y | 8 | 630 | 188 | 0.792 (0.772–0.810) | 0.649 (0.618–0.679) |
| ETHOS | HbA1c ≤6% to E11, 5 y | 16 | 630 | 188 | 0.828 (0.811–0.844) | 0.702 (0.677–0.728) |
| ETHOS | HbA1c ≤6% to E11, 5 y | 32 | 630 | 188 | 0.852 (0.842–0.862) | 0.740 (0.725–0.754) |
| ETHOS | HbA1c ≤6% to E11, 5 y | 64 | 630 | 188 | 0.863 (0.853–0.872) | 0.755 (0.738–0.773) |

| Implementation | Task | $K$ | Test $n$ | Positive $n$ | AUROC (95% CI) | AP (95% CI) |
|---|---|---|---|---|---|---|
| ETHOS | HbA1c ≤6% to E11, 5 y | 128 | 630 | 188 | 0.885 (0.881–0.888) | 0.795 (0.788–0.802) |
| ETHOS | HbA1c >6 to <7.5% to E11, 5 y | 8 | 517 | 394 | 0.700 (0.657–0.738) | 0.874 (0.851–0.894) |
| ETHOS | HbA1c >6 to <7.5% to E11, 5 y | 16 | 517 | 394 | 0.768 (0.749–0.785) | 0.911 (0.902–0.921) |
| ETHOS | HbA1c >6 to <7.5% to E11, 5 y | 32 | 517 | 394 | 0.781 (0.765–0.797) | 0.919 (0.911–0.927) |
| ETHOS | HbA1c >6 to <7.5% to E11, 5 y | 64 | 517 | 394 | 0.797 (0.786–0.806) | 0.927 (0.923–0.931) |
| ETHOS | HbA1c >6 to <7.5% to E11, 5 y | 128 | 517 | 394 | Not estimable | Not estimable |
| ETHOS | HbA1c ≥7.5% to E11, 2 y | 8 | 367 | 303 | 0.701 (0.669–0.729) | 0.911 (0.898–0.923) |
| ETHOS | HbA1c ≥7.5% to E11, 2 y | 16 | 367 | 303 | 0.742 (0.719–0.765) | 0.930 (0.923–0.938) |
| ETHOS | HbA1c ≥7.5% to E11, 2 y | 32 | 367 | 303 | 0.775 (0.762–0.786) | 0.940 (0.936–0.944) |
| ETHOS | HbA1c ≥7.5% to E11, 2 y | 64 | 367 | 303 | 0.807 (0.799–0.816) | 0.950 (0.948–0.953) |
| ETHOS | HbA1c ≥7.5% to E11, 2 y | 128 | 367 | 303 | Not estimable | Not estimable |
| ETHOS | E11 to A10B, 2 y | 8 | 479 | 117 | 0.537 (0.525–0.548) | 0.281 (0.271–0.290) |
| ETHOS | E11 to A10B, 2 y | 16 | 479 | 117 | 0.557 (0.540–0.574) | 0.292 (0.278–0.308) |
| ETHOS | E11 to A10B, 2 y | 32 | 479 | 117 | 0.574 (0.561–0.589) | 0.307 (0.295–0.319) |
| ETHOS | E11 to A10B, 2 y | 64 | 479 | 117 | 0.578 (0.565–0.591) | 0.315 (0.302–0.329) |
| ETHOS | E11 to A10B, 2 y | 128 | 479 | 117 | Not estimable | Not estimable |
| ETHOS | E11 to A10A, 2 y | 8 | 178 | 89 | 0.541 (0.518–0.561) | 0.560 (0.539–0.579) |
| ETHOS | E11 to A10A, 2 y | 16 | 178 | 89 | 0.600 (0.582–0.617) | 0.613 (0.595–0.630) |
| ETHOS | E11 to A10A, 2 y | 32 | 178 | 89 | 0.593 (0.575–0.611) | 0.606 (0.587–0.625) |
| ETHOS | E11 to A10A, 2 y | 64 | 178 | 89 | 0.606 (0.596–0.616) | 0.601 (0.592–0.610) |
| ETHOS | E11 to A10A, 2 y | 128 | 178 | 89 | Not estimable | Not estimable |
| ETHOS | E11 to A10A, 7 y | 8 | 146 | 130 | Not estimable | Not estimable |
| ETHOS | E11 to A10A, 7 y | 16 | 146 | 130 | Not estimable | Not estimable |
| ETHOS | E11 to A10A, 7 y | 32 | 146 | 130 | Not estimable | Not estimable |
| ETHOS | E11 to A10A, 7 y | 64 | 146 | 130 | Not estimable | Not estimable |
| ETHOS | E11 to A10A, 7 y | 128 | 146 | 130 | Not estimable | Not estimable |
| ETHOS | E11 to N18, 2 y | 8 | 804 | 405 | 0.827 (0.807–0.846) | 0.831 (0.811–0.849) |
| ETHOS | E11 to N18, 2 y | 16 | 804 | 405 | 0.851 (0.825–0.874) | 0.857 (0.830–0.880) |
| ETHOS | E11 to N18, 2 y | 32 | 804 | 405 | 0.894 (0.887–0.901) | 0.898 (0.888–0.907) |
| ETHOS | E11 to N18, 2 y | 64 | 804 | 405 | 0.903 (0.895–0.910) | 0.910 (0.903–0.916) |
| ETHOS | E11 to N18, 2 y | 128 | 804 | 405 | 0.914 (0.908–0.919) | 0.921 (0.915–0.926) |
| ETHOS | E11 to N18, 10 y | 8 | 500 | 475 | 0.912 (0.897–0.925) | 0.994 (0.993–0.995) |
| ETHOS | E11 to N18, 10 y | 16 | 500 | 475 | 0.938 (0.930–0.944) | 0.996 (0.995–0.997) |
| ETHOS | E11 to N18, 10 y | 32 | 500 | 475 | Not estimable | Not estimable |
| ETHOS | E11 to N18, 10 y | 64 | 500 | 475 | Not estimable | Not estimable |
| ETHOS | E11 to N18, 10 y | 128 | 500 | 475 | Not estimable | Not estimable |
| ETHOS | E11 to H36, 5 y | 8 | 279 | 0 | Not estimable | Not estimable |
| ETHOS | E11 to H36, 5 y | 16 | 279 | 0 | Not estimable | Not estimable |
| ETHOS | E11 to H36, 5 y | 32 | 279 | 0 | Not estimable | Not estimable |
| ETHOS | E11 to H36, 5 y | 64 | 279 | 0 | Not estimable | Not estimable |
| ETHOS | E11 to H36, 5 y | 128 | 279 | 0 | Not estimable | Not estimable |
| ETHOS | E11 to G63/G62.9, 5 y | 8 | 293 | 15 | Not estimable | Not estimable |
| ETHOS | E11 to G63/G62.9, 5 y | 16 | 293 | 15 | Not estimable | Not estimable |
| ETHOS | E11 to G63/G62.9, 5 y | 32 | 293 | 15 | Not estimable | Not estimable |
| ETHOS | E11 to G63/G62.9, 5 y | 64 | 293 | 15 | Not estimable | Not estimable |
| ETHOS | E11 to G63/G62.9, 5 y | 128 | 293 | 15 | Not estimable | Not estimable |
| ETHOS | E11 to I25/I50/I63, 5 y | 8 | 906 | 760 | 0.790 (0.761–0.816) | 0.946 (0.936–0.955) |
| ETHOS | E11 to I25/I50/I63, 5 y | 16 | 906 | 760 | 0.799 (0.779–0.817) | 0.949 (0.942–0.955) |
| ETHOS | E11 to I25/I50/I63, 5 y | 32 | 906 | 760 | 0.847 (0.837–0.857) | 0.965 (0.962–0.968) |
| ETHOS | E11 to I25/I50/I63, 5 y | 64 | 906 | 760 | 0.837 (0.823–0.850) | 0.963 (0.958–0.966) |
| ETHOS | E11 to I25/I50/I63, 5 y | 128 | 906 | 760 | 0.861 (0.855–0.867) | 0.970 (0.968–0.972) |
| ETHOS | E11 to death, 10 y | 8 | 484 | 439 | 0.840 (0.825–0.856) | 0.979 (0.977–0.982) |
| ETHOS | E11 to death, 10 y | 16 | 484 | 439 | 0.856 (0.840–0.872) | 0.981 (0.979–0.984) |
| ETHOS | E11 to death, 10 y | 32 | 484 | 439 | 0.886 (0.878–0.894) | 0.986 (0.985–0.988) |
| ETHOS | E11 to death, 10 y | 64 | 484 | 439 | Not estimable | Not estimable |
| ETHOS | E11 to death, 10 y | 128 | 484 | 439 | Not estimable | Not estimable |
| Delphi demo | HbA1c ≤6% to E11, 5 y | 8 | 0 | 0 | Not estimable | Not estimable |
| Delphi demo | HbA1c ≤6% to E11, 5 y | 16 | 0 | 0 | Not estimable | Not estimable |
| Delphi demo | HbA1c ≤6% to E11, 5 y | 32 | 0 | 0 | Not estimable | Not estimable |
| Delphi demo | HbA1c ≤6% to E11, 5 y | 64 | 0 | 0 | Not estimable | Not estimable |
| Delphi demo | HbA1c ≤6% to E11, 5 y | 128 | 0 | 0 | Not estimable | Not estimable |
| Delphi demo | HbA1c >6 to <7.5% to E11, 5 y | 8 | 0 | 0 | Not estimable | Not estimable |
| Delphi demo | HbA1c >6 to <7.5% to E11, 5 y | 16 | 0 | 0 | Not estimable | Not estimable |
| Delphi demo | HbA1c >6 to <7.5% to E11, 5 y | 32 | 0 | 0 | Not estimable | Not estimable |
| Delphi demo | HbA1c >6 to <7.5% to E11, 5 y | 64 | 0 | 0 | Not estimable | Not estimable |
| Delphi demo | HbA1c >6 to <7.5% to E11, 5 y | 128 | 0 | 0 | Not estimable | Not estimable |
| Delphi demo | HbA1c ≥7.5% to E11, 2 y | 8 | 0 | 0 | Not estimable | Not estimable |
| Delphi demo | HbA1c ≥7.5% to E11, 2 y | 16 | 0 | 0 | Not estimable | Not estimable |
| Delphi demo | HbA1c ≥7.5% to E11, 2 y | 32 | 0 | 0 | Not estimable | Not estimable |
| Delphi demo | HbA1c ≥7.5% to E11, 2 y | 64 | 0 | 0 | Not estimable | Not estimable |

| Implementation | Task | $K$ | Test $n$ | Positive $n$ | AUROC (95% CI) | AP (95% CI) |
|---|---|---|---|---|---|---|
| Delphi demo | HbA1c ≥7.5% to E11, 2 y | 128 | 0 | 0 | Not estimable | Not estimable |
| Delphi demo | E11 to A10B, 2 y | 8 | 0 | 0 | Not estimable | Not estimable |
| Delphi demo | E11 to A10B, 2 y | 16 | 0 | 0 | Not estimable | Not estimable |
| Delphi demo | E11 to A10B, 2 y | 32 | 0 | 0 | Not estimable | Not estimable |
| Delphi demo | E11 to A10B, 2 y | 64 | 0 | 0 | Not estimable | Not estimable |
| Delphi demo | E11 to A10B, 2 y | 128 | 0 | 0 | Not estimable | Not estimable |
| Delphi demo | E11 to A10A, 2 y | 8 | 0 | 0 | Not estimable | Not estimable |
| Delphi demo | E11 to A10A, 2 y | 16 | 0 | 0 | Not estimable | Not estimable |
| Delphi demo | E11 to A10A, 2 y | 32 | 0 | 0 | Not estimable | Not estimable |
| Delphi demo | E11 to A10A, 2 y | 64 | 0 | 0 | Not estimable | Not estimable |
| Delphi demo | E11 to A10A, 2 y | 128 | 0 | 0 | Not estimable | Not estimable |
| Delphi demo | E11 to A10A, 7 y | 8 | 0 | 0 | Not estimable | Not estimable |
| Delphi demo | E11 to A10A, 7 y | 16 | 0 | 0 | Not estimable | Not estimable |
| Delphi demo | E11 to A10A, 7 y | 32 | 0 | 0 | Not estimable | Not estimable |
| Delphi demo | E11 to A10A, 7 y | 64 | 0 | 0 | Not estimable | Not estimable |
| Delphi demo | E11 to A10A, 7 y | 128 | 0 | 0 | Not estimable | Not estimable |
| Delphi demo | E11 to N18, 2 y | 8 | 387 | 5 | Not estimable | Not estimable |
| Delphi demo | E11 to N18, 2 y | 16 | 387 | 5 | Not estimable | Not estimable |
| Delphi demo | E11 to N18, 2 y | 32 | 387 | 5 | Not estimable | Not estimable |
| Delphi demo | E11 to N18, 2 y | 64 | 387 | 5 | Not estimable | Not estimable |
| Delphi demo | E11 to N18, 2 y | 128 | 387 | 5 | Not estimable | Not estimable |
| Delphi demo | E11 to N18, 10 y | 8 | 223 | 26 | 0.725 (0.696–0.753) | 0.309 (0.272–0.349) |
| Delphi demo | E11 to N18, 10 y | 16 | 223 | 26 | 0.728 (0.698–0.759) | 0.306 (0.266–0.353) |
| Delphi demo | E11 to N18, 10 y | 32 | 223 | 26 | Not estimable | Not estimable |
| Delphi demo | E11 to N18, 10 y | 64 | 223 | 26 | Not estimable | Not estimable |
| Delphi demo | E11 to N18, 10 y | 128 | 223 | 26 | Not estimable | Not estimable |
| Delphi demo | E11 to H36, 5 y | 8 | 334 | 6 | Not estimable | Not estimable |
| Delphi demo | E11 to H36, 5 y | 16 | 334 | 6 | Not estimable | Not estimable |
| Delphi demo | E11 to H36, 5 y | 32 | 334 | 6 | Not estimable | Not estimable |
| Delphi demo | E11 to H36, 5 y | 64 | 334 | 6 | Not estimable | Not estimable |
| Delphi demo | E11 to H36, 5 y | 128 | 334 | 6 | Not estimable | Not estimable |
| Delphi demo | E11 to G63/G62.9, 5 y | 8 | 0 | 0 | Not estimable | Not estimable |
| Delphi demo | E11 to G63/G62.9, 5 y | 16 | 0 | 0 | Not estimable | Not estimable |
| Delphi demo | E11 to G63/G62.9, 5 y | 32 | 0 | 0 | Not estimable | Not estimable |
| Delphi demo | E11 to G63/G62.9, 5 y | 64 | 0 | 0 | Not estimable | Not estimable |
| Delphi demo | E11 to G63/G62.9, 5 y | 128 | 0 | 0 | Not estimable | Not estimable |
| Delphi demo | E11 to I25/I50/I63, 5 y | 8 | 319 | 38 | 0.620 (0.591–0.647) | 0.216 (0.186–0.252) |
| Delphi demo | E11 to I25/I50/I63, 5 y | 16 | 319 | 38 | 0.605 (0.582–0.630) | 0.212 (0.189–0.236) |
| Delphi demo | E11 to I25/I50/I63, 5 y | 32 | 319 | 38 | 0.651 (0.634–0.667) | 0.259 (0.238–0.278) |
| Delphi demo | E11 to I25/I50/I63, 5 y | 64 | 319 | 38 | Not estimable | Not estimable |
| Delphi demo | E11 to I25/I50/I63, 5 y | 128 | 319 | 38 | Not estimable | Not estimable |
| Delphi demo | E11 to death, 10 y | 8 | 313 | 97 | 0.886 (0.866–0.906) | 0.820 (0.785–0.853) |
| Delphi demo | E11 to death, 10 y | 16 | 313 | 97 | 0.910 (0.896–0.923) | 0.855 (0.835–0.873) |
| Delphi demo | E11 to death, 10 y | 32 | 313 | 97 | 0.930 (0.922–0.938) | 0.891 (0.877–0.905) |
| Delphi demo | E11 to death, 10 y | 64 | 313 | 97 | 0.938 (0.934–0.942) | 0.907 (0.902–0.913) |
| Delphi demo | E11 to death, 10 y | 128 | 313 | 97 | Not estimable | Not estimable |
| CEHR-GPT demo | HbA1c ≤6% to E11, 5 y | 8 | 0 | 0 | Not estimable | Not estimable |
| CEHR-GPT demo | HbA1c ≤6% to E11, 5 y | 16 | 0 | 0 | Not estimable | Not estimable |
| CEHR-GPT demo | HbA1c ≤6% to E11, 5 y | 32 | 0 | 0 | Not estimable | Not estimable |
| CEHR-GPT demo | HbA1c ≤6% to E11, 5 y | 64 | 0 | 0 | Not estimable | Not estimable |
| CEHR-GPT demo | HbA1c ≤6% to E11, 5 y | 128 | 0 | 0 | Not estimable | Not estimable |
| CEHR-GPT demo | HbA1c >6 to <7.5% to E11, 5 y | 8 | 0 | 0 | Not estimable | Not estimable |
| CEHR-GPT demo | HbA1c >6 to <7.5% to E11, 5 y | 16 | 0 | 0 | Not estimable | Not estimable |
| CEHR-GPT demo | HbA1c >6 to <7.5% to E11, 5 y | 32 | 0 | 0 | Not estimable | Not estimable |
| CEHR-GPT demo | HbA1c >6 to <7.5% to E11, 5 y | 64 | 0 | 0 | Not estimable | Not estimable |
| CEHR-GPT demo | HbA1c >6 to <7.5% to E11, 5 y | 128 | 0 | 0 | Not estimable | Not estimable |
| CEHR-GPT demo | HbA1c ≥7.5% to E11, 2 y | 8 | 0 | 0 | Not estimable | Not estimable |
| CEHR-GPT demo | HbA1c ≥7.5% to E11, 2 y | 16 | 0 | 0 | Not estimable | Not estimable |

| Implementation | Task | $K$ | Test $n$ | Positive $n$ | AUROC (95% CI) | AP (95% CI) |
|---|---|---|---|---|---|---|
| CEHR-GPT demo | HbA1c ≥7.5% to E11, 2 y | 32 | 0 | 0 | Not estimable | Not estimable |
| CEHR-GPT demo | HbA1c ≥7.5% to E11, 2 y | 64 | 0 | 0 | Not estimable | Not estimable |
| CEHR-GPT demo | HbA1c ≥7.5% to E11, 2 y | 128 | 0 | 0 | Not estimable | Not estimable |
| CEHR-GPT demo | E11 to A10B, 2 y | 8 | 1,240 | 336 | 0.534 (0.516–0.552) | 0.304 (0.292–0.317) |
| CEHR-GPT demo | E11 to A10B, 2 y | 16 | 1,240 | 336 | 0.549 (0.529–0.568) | 0.314 (0.299–0.329) |
| CEHR-GPT demo | E11 to A10B, 2 y | 32 | 1,240 | 336 | 0.558 (0.544–0.572) | 0.321 (0.310–0.332) |
| CEHR-GPT demo | E11 to A10B, 2 y | 64 | 1,240 | 336 | 0.564 (0.551–0.575) | 0.325 (0.316–0.333) |
| CEHR-GPT demo | E11 to A10B, 2 y | 128 | 1,240 | 336 | 0.588 (0.578–0.597) | 0.345 (0.336–0.354) |
| CEHR-GPT demo | E11 to A10A, 2 y | 8 | 1,252 | 86 | 0.494 (0.470–0.521) | 0.073 (0.067–0.080) |
| CEHR-GPT demo | E11 to A10A, 2 y | 16 | 1,252 | 86 | 0.483 (0.464–0.501) | 0.074 (0.069–0.080) |
| CEHR-GPT demo | E11 to A10A, 2 y | 32 | 1,252 | 86 | 0.493 (0.482–0.504) | 0.072 (0.070–0.075) |
| CEHR-GPT demo | E11 to A10A, 2 y | 64 | 1,252 | 86 | Not estimable | Not estimable |
| CEHR-GPT demo | E11 to A10A, 2 y | 128 | 1,252 | 86 | Not estimable | Not estimable |
| CEHR-GPT demo | E11 to A10A, 7 y | 8 | 1,095 | 280 | 0.496 (0.479–0.512) | 0.263 (0.252–0.273) |
| CEHR-GPT demo | E11 to A10A, 7 y | 16 | 1,095 | 280 | 0.499 (0.481–0.517) | 0.269 (0.257–0.282) |
| CEHR-GPT demo | E11 to A10A, 7 y | 32 | 1,095 | 280 | 0.507 (0.494–0.520) | 0.271 (0.262–0.280) |
| CEHR-GPT demo | E11 to A10A, 7 y | 64 | 1,095 | 280 | 0.508 (0.500–0.516) | 0.272 (0.266–0.278) |
| CEHR-GPT demo | E11 to A10A, 7 y | 128 | 1,095 | 280 | 0.510 (0.502–0.519) | 0.275 (0.268–0.282) |
| CEHR-GPT demo | E11 to N18, 2 y | 8 | 1,253 | 110 | 0.502 (0.480–0.525) | 0.096 (0.089–0.104) |
| CEHR-GPT demo | E11 to N18, 2 y | 16 | 1,253 | 110 | 0.484 (0.459–0.505) | 0.091 (0.084–0.099) |
| CEHR-GPT demo | E11 to N18, 2 y | 32 | 1,253 | 110 | 0.496 (0.481–0.512) | 0.096 (0.092–0.101) |
| CEHR-GPT demo | E11 to N18, 2 y | 64 | 1,253 | 110 | Not estimable | Not estimable |
| CEHR-GPT demo | E11 to N18, 2 y | 128 | 1,253 | 110 | Not estimable | Not estimable |
| CEHR-GPT demo | E11 to N18, 10 y | 8 | 1,017 | 358 | 0.520 (0.507–0.531) | 0.396 (0.378–0.414) |
| CEHR-GPT demo | E11 to N18, 10 y | 16 | 1,017 | 358 | 0.518 (0.506–0.530) | 0.387 (0.373–0.402) |
| CEHR-GPT demo | E11 to N18, 10 y | 32 | 1,017 | 358 | 0.530 (0.517–0.543) | 0.396 (0.382–0.410) |
| CEHR-GPT demo | E11 to N18, 10 y | 64 | 1,017 | 358 | 0.534 (0.526–0.543) | 0.410 (0.398–0.423) |
| CEHR-GPT demo | E11 to N18, 10 y | 128 | 1,017 | 358 | 0.548 (0.539–0.556) | 0.440 (0.433–0.447) |
| CEHR-GPT demo | E11 to H36, 5 y | 8 | 0 | 0 | Not estimable | Not estimable |
| CEHR-GPT demo | E11 to H36, 5 y | 16 | 0 | 0 | Not estimable | Not estimable |
| CEHR-GPT demo | E11 to H36, 5 y | 32 | 0 | 0 | Not estimable | Not estimable |
| CEHR-GPT demo | E11 to H36, 5 y | 64 | 0 | 0 | Not estimable | Not estimable |
| CEHR-GPT demo | E11 to H36, 5 y | 128 | 0 | 0 | Not estimable | Not estimable |
| CEHR-GPT demo | E11 to G63/G62.9, 5 y | 8 | 0 | 0 | Not estimable | Not estimable |
| CEHR-GPT demo | E11 to G63/G62.9, 5 y | 16 | 0 | 0 | Not estimable | Not estimable |

| Implementation | Task | $K$ | Test $n$ | Positive $n$ | AUROC (95% CI) | AP (95% CI) |
|---|---|---|---|---|---|---|
| CEHR-GPT demo | E11 to G63/G62.9, 5 y | 32 | 0 | 0 | Not estimable | Not estimable |
| CEHR-GPT demo | E11 to G63/G62.9, 5 y | 64 | 0 | 0 | Not estimable | Not estimable |
| CEHR-GPT demo | E11 to G63/G62.9, 5 y | 128 | 0 | 0 | Not estimable | Not estimable |
| CEHR-GPT demo | E11 to I25/I50/I63, 5 y | 8 | 1,131 | 13 | Not estimable | Not estimable |
| CEHR-GPT demo | E11 to I25/I50/I63, 5 y | 16 | 1,131 | 13 | Not estimable | Not estimable |
| CEHR-GPT demo | E11 to I25/I50/I63, 5 y | 32 | 1,131 | 13 | Not estimable | Not estimable |
| CEHR-GPT demo | E11 to I25/I50/I63, 5 y | 64 | 1,131 | 13 | Not estimable | Not estimable |
| CEHR-GPT demo | E11 to I25/I50/I63, 5 y | 128 | 1,131 | 13 | Not estimable | Not estimable |
| CEHR-GPT demo | E11 to death, 10 y | 8 | 0 | 0 | Not estimable | Not estimable |
| CEHR-GPT demo | E11 to death, 10 y | 16 | 0 | 0 | Not estimable | Not estimable |
| CEHR-GPT demo | E11 to death, 10 y | 32 | 0 | 0 | Not estimable | Not estimable |
| CEHR-GPT demo | E11 to death, 10 y | 64 | 0 | 0 | Not estimable | Not estimable |
| CEHR-GPT demo | E11 to death, 10 y | 128 | 0 | 0 | Not estimable | Not estimable |
